# Unrestricted Versus Regulated Open Data Governance: A Bibliometric Comparison of SARS-CoV-2 Nucleotide Sequence Databases

**DOI:** 10.1101/2023.05.13.540634

**Authors:** Nathanael Sheehan, Federico Botta, Sabina Leonelli

**Affiliations:** University of Exeter, College of Engineering, Mathematics, and Physical Sciences; University of Exeter, Egenis, the Centre for the Study of Life Sciences

**Author notes:** For the purpose of open access, the author has applied a Creative Commons Attribution (CC BY) licence to any Author Accepted Manuscript version arising from this submission. Funding Statement: This project has received funding from the European Research Council (ERC) under the European Union’s Horison 2020 research and innovation programme (grant agreement No. 101001145). This paper reflects only the authors’ views and the Commission /Agency is not responsible for any use that may be made of the information it contains. Author Contribution statement: N.S and S.L conceived the presented idea. N.S, S.L and F.B contributed to the design and implementation of the research. N.S and F.B performed the data analysis and N.S and S.L wrote the manuscript. Conflict of Interest: The authors declare that they have no conflict of interest.

**Keywords:** *Covid-19*, *Genomic Data Sharing*, *Data Infrastructures*, *Data Governance*, *Open Science*, *Metascience*

## Abstract

Two distinct modes of data governance have emerged in accessing and reusing viral data pertaining to COVID-19: an unrestricted model, espoused by data repositories part of the International Nucleotide Sequence Database Collaboration and a regulated model promoted by the Global Initiative on Sharing All Influenza data. In this paper, we focus on publications mentioning either infrastructure in the period between January 2020 and January 2023, thus capturing a period of acute response to the COVID-19 pandemic. Through a variety of bibliometric and network science methods, we compare the extent to which either data infrastructure facilitated collaboration from different countries around the globe to understand how data reuse can enhance forms of diversity between institutions, countries, and funding groups. Our findings reveal disparities in representation and usage between the two data infrastructures. We conclude that both approaches offer useful lessons, with the unrestricted model providing insights into complex data linkage and the regulated model demonstrating the importance of global representation.

## 1 Introduction

To date no other scientific data has been shared as widely as the genetic information on the SARS-CoV-2 virus responsible for the recent pandemic. This phenomenon has led some scholars, such as Leach (2021), to argue that alongside the COVID-19 pandemic, we are witnessing another kind of outbreak, where the act of sharing data has evolved into active participation in Open Science, giving rise to an “open data pandemic”. While the notions of Open Data (OD) and Open Science (OS) have been topics of discourse among scientists, philosophers, and policymakers for well over two decades, they have gained unprecedented prominence during the COVID-19 crisis both within and outside of science. This surge in interest and application reflects a broader call for increased transparency, inclusivity, and accountability in data-centric research (Burgelman 2019). This push towards openness has led to numerous national and international policies implementing infrastructures, principles, and resources in a top-down fashion, which is not always aligned with what specific groups of actors understand responsible OD or OS to be (Leonelli 2023).

This study focuses on the contrasting repertoires (Ankeny and Leonelli 2016) of data governance that have arisen in accessing and reusing viral data concerning SARS-CoV-2. So far at least two distinct models have emerged: the unrestricted model endorsed by data repositories within the International Nucleotide Sequence Database Collaboration (INSDC RRID:SCR_011967) and the regulated model promoted by the Global Initiative on Sharing All Influenza data (GISAID RRID:SCR_018251).^1^ The former encourages free access to data with no constraints, emphasising interpretable and rapid dissemination, while the latter maintains free data access but implements certain constraints and rules around data usage to address issues of credit attribution and exploitation. A notable global health concern amidst these models is the challenge of equity and inclusion within the research landscape. The barriers and limitations that researchers from low-resourced environments and less-visible research locations face, including problems with receiving due credit for their contributions and participating in subsequent research and development (Bezuidenhout et al 2017), have prompted the development of governance strategies along the lines implemented by GISAID (Khan 2017). Yet, in the midst of these endeavours to ensure inclusivity and fairness, GISAID’s regulated data model became a point of contention, drawing repeated criticisms from the INDSC during the height of the pandemic for not being open enough and thereby not supporting response efforts as well as the emergency required (EBI 2021; Enserink and Cohen 2023).

The study is presented as follows: first we provide an account of the two data infrastructures and their principles of sound data management. We then go on to compare the characteristics of bibliometric indicators, access patterns, publishers, key terms, viral variants, research collaborations, and funding dynamics for GISAID and each of the repositories that make up the INSDC – The European Nucleotide Archive (ENA), National Center for Biotechnology Information (NCBI) and the DNA Data Bank of Japan (DDBJ). Our analysis is grounded on data from Digital Science’s Dimensions.ai literature database which mentions either repository in the period between January 2020 and January 2023. We conclude with a reflection on what these two modes of data governance have achieved in terms of their representativeness and interpretations of openness.

## 2 Background: Two models of Governance for Covid-19 Data

### 2.1 GISAID: A Regulated Access Model of Open Data Governance

The GISAID Epi-Flu database was launched in 2008, on the anniversary of the Spanish influenza, to foster the sharing of influenza genomic data securely and responsibly. Data sharing was immediately conceptualised not as straightforward ‘opening up’ of the data by placing them online without restrictions to access and re-use, but rather as an alternative to the public sharing model, whereby users agree to authenticate their academic identity and not to republish or link GISAID genomes without permission from the data producer. GISAID acts as a mediator and enforcer of such rules, granting access solely to users who credibly profess to adopt them, and thereby acting as a guarantor for the effectiveness of the sharing agreement entered by data contributors. This arrangement stems from the recognition that some researchers – commonly located in low-resourced environments – are reluctant to share data due to fears of better-equipped researchers building on such work without due acknowledgment (Elbe and Buckland-Merrett 2017; Bezuidenhout and Chakauya 2018). This model proved successful in facilitating better credit attribution to contributing scientists in relation to influenza research, and since its launch GISAID has played an essential role supporting data sharing among the WHO Collaborating Centers and National influenza Centers in response to the bi-annual influenza vaccine virus recommendations by the WHO Global influenza Surveillance and Response System (GISRS). It is no surprise therefore that GISAID was swiftly redeployed, in early 2020, to include SARS-CoV-2 data through the EpiCov database, which stores, analyses and builds evolutionary trees of SARS CoV-2 genome sequences and hosts several daily updates of visualisations (Khare et al 2021). GISAID is now the leading open access database for SARS-CoV-2, with over 15 million genomes sequenced by February 2023.The Epi-CoV database provides eleven tools to explore SARS-CoV-2 sequence data, within in these include Audacity – global phylogeny of hCoV-19 as a downloadable newick tree file -, CoVizue-near real-time visualisation of SARS-CoV-2 genetic variation – as well as a collection of thirty-two analysis figures updated daily.

GISAID has played a key role in supporting the identification and study of variant evolution, lineages, and spread in real-time during the first three years of the pandemic; and at the time of writing it still features as key data provider for a wide variety of consortia, initiatives and projects devoted to the analysis of COVID-19 variants of interest (some of which listed on this page: https://gisaid.org/collaborations/enabled-by-hcov-19-data-from-gisaid/). Accordingly, GISAID is funded by a wide consortium of public and private bodies, including the Federal Republic of Germany, who first backed the project at its main site in Geneva, as well as public-health and academic institutions in Argentina, Brazil, China, Republic of the Congo, Ethiopia, Indonesia, Malaysia, Russia, Senegal, Singapore, South Africa and many other countries, as well as several donors and partners garnered under the label of “Friends of GISAID”.

The GISAID model has fostered information exchange among groups that differ considerably in their geo-political locations, funding levels, material resources and social characteristics, thereby expanding the range of data sources shared online (Shu and McCauley 2017). At the same time, GISAID has been frequently scrutinised as limiting the extent to which data can be accessed and linked, thereby negatively affect the insight, pace and breadth of future research – leading to the backlash by hundreds of leading researchers concerned about the urgency of an effective pandemic response (ENA 2021). During the height of the pandemic, questions were also raised regarding the quality and integrity of metadata coming from the GISAID platform (Gozashti and Corbett-Detig 2021), as well as epistemic importance being placed on the lag in submissions times at the global scale (Kalia et al. 2021). Some scientists called for a complete opening of genomic data sharing for SARS-CoV-2 (Van Noorden 2021), stating that GISAID’s policy may be “open data”, but it does not make the data easily shareable or useable (Yehudi et al 2022). These critiques have run alongside controversy around who retains ownership of the data stored in GISAID, and how reliably its governance is actually managed. During past viral outbreaks, GISAID has been involved in legal disputes between the Swiss Institute of Bioinformatics (SIB) over monetary funds of the infrastructure (Greenemeier 2009), events which led to a spokeswoman at GISAID asserting that the SIB had misappropriated the database on grounds of data ownership (Butler 2009). Recent allegations also emerged around GISAID refusing to share data with researchers who, despite complying with its policies, had been critical of them (Enserink and Cohen 2023). Though worth mentioning here given their heated and prominent nature, this paper does not concern itself with these debates, focusing instead on the ways in which GISAID data have been accessed and used.

### 2.2 INSDC: An Unrestricted Access Model of Open Data Governance

The fully open model promoted by the International Nucleotide Sequence Database Collaboration (INSDC) has its origins in the first nucleotide sequence database, dating back to the development of the Data Library at the European Molecular Biology Laboratory (EMBL RRID:SCR_004473) in 1982 Heidelberg, Germany. Soon after its development, the database collaborated with GenBank (RRID:SCR_002760) at the Los Alamos Science Laboratory, USA (Arita et al., 2021a) and finally with the DNA data bank of Japan (DDBJ RRID:SCR_002359) in 1987 (Fukuda et al., 2021), resulting in what Bernasconi et al (2020) call a “political integration of sequences”.

By 2002, the governing board of the INSDC had published a data sharing policy which permitted free and unrestricted access to all data in the triad of repositories. The policy makes all data records immediately available to all users, commercial sectors included, without any licensing stipulations, access restrictions, or monetary charges (Karsch-Mizrachi, 2018). Internally, all data shared with either node of the INSDC would be mirrored into the other repositories. The INSDC’s policies of unrestrictive data sharing follow in the steps of the Bermuda Principles of 1996, which established the norm of full and immediate data sharing for genomic information within biology (Maxon et al 2018); at the same time, they precede the institutionalization of the Open Access Movement in academic publishing and institutes such as Open Data Institute (ODI RRID:SCR_021681), effectively making it an avant-garde institution in the broader context of Open Science (Arita, 2021b).

After four decades of operation, INSDC has displayed a remarkable constancy, with superficial changes largely limited to institutional rebranding. For example, the EMBL Data Library has metamorphosed into the European Nucleotide Archive (ENA RRID:SCR_006515), now under the aegis of the European Bioinformatics Institute in Cambridge Hinxton, while GenBank falls under the purview of the National Center for Biotechnology Information (NCBI RRID:SCR_006472) in Bethesda, USA. Amidst the urgency of the COVID-19 pandemic, the INSDC’s well-established data infrastructures offered critical support for data submission, harmonisation, and the ability to publicly access sequence data with a range of new and old tools and portals. For the NCBI these included: Genebank for annotated sequences of RNA and DNA data; RefSeq for tasks ranging from genome annotations to mutation and polymorphism evaluations, as well as the development of the NCBI virus data dashboard and the NCBI Covid Hub (Berasconi et al., 2021).Additionally, in April 2020, the European Commission, through the auspices of the European Molecular Biology Laboratory-European Bioinformatics Institute (EMBL-EBI) and Elixir (a twenty-three country node network in Europe dedicated to openly sharing life science data) launched another platform for sharing scientific information of relevance to the biological study of COVID-19: the COVID-19 Data Portal (CV19-DP RRID:SCR_018337) (Harrison et al 2021). This data infrastructure hosts a diverse array of genetic, epidemiological, socio-economic and literature data, encouraging data linkage and cross-analysis (Saravanan et al. 2022). Notably, CV19-DP features a modular design that allows for the rapid development of nation-specific customized interfaces; examples include versions for Spain, Sweden and Poland. However, the primary portal remains the principal access point. One of the overarching aims of CV19-DP is to expedite scientific research by enhancing data interoperability across various bioinformatics platforms, such as ENA, UniProt (RRID:SCR_002380), PDBe (RRID:SCR_004312), EMDB (RRID:SCR_003207), Expression Atlas (RRID:SCR_007989), and Europe PMC (RRID:SCR_005901). To achieve this, the portal deploys a high-level Application Programming Interface (API) and supports direct bulk downloads, ensuring minimal user tracking for transparent and efficient data dissemination.

CV19-DP provides two primary data visualization tools. The first is an open-source phylogenetic tree that displays COVID-19 sequences, constituting 98% of the reference SARS-CoV-2 genome, including PANGO lineages stratified by World Health Organization (WHO) regions. The second is CoVeo, a proprietary browser that performs systematic analysis of raw sequence data, providing visual and summary analyses for various regions, particularly focused on Variants of Concern (VOCs) and Variants of Interest (VOIs) (Rahman et al., 2023).The interface of CV19-DP was later adapted for the Pathogens Hub by the European Nucleotide Archive (ENA), launched in July 2023. This hub is a curated repository founded on the UK’s Health and Safety Executive’s list of approved biological agents, as well as the WHO’s global priority pathogens list. Although other interfaces exist within the INDSC such as NCBI Virus (RRID:SCR_018253), and the EBI SARS-CoV-2 data hub, the CV19-DP has assumed a significant role. It notably hosted an open letter advocating for the unreserved sharing of SARS-CoV-2 resources and urging submissions to one of the INSDC databases. While the letter did not explicitly name GISAID, it was widely interpreted as a critique of GISAID’s data governance and vision.

### 2.3 Principles of Sound Data Management

For CV19-DP, and the rest of the cadre in the INSDC, sound data management is predicated on the FAIR data principles, which espouse the findability, accessibility, interoperability, and reusability of research data (Wilkinson et al., 2016). This was justified by the ENA with regards to the publication of CV19-DP where they state

> “…unrestricted access to data plays a critical role in the rapid coronavirus research necessary to respond to this global health crisis…. Ensuring open science and unrestricted international collaborations is of key importance, and it is recognised that these datasets must be shared openly and meet FAIR standards” (Harrison et al 2021).

The FAIR principles are also found on the NCBI Covid-19 Data Submissions page, where they state a benefit of submitting data to them is to *“follow FAIR data-sharing principles”.* The principles of FAIR have been embraced by the a number of different research areas, this is demonstrated by the growing body of literature on FAIR data sharing (Stall et al. 2019; Wise et al.2019; Bezuidenhout 2020;Goble et al. 2021;Leonelli 2021;European Commission 2022) and the cross fertilization to principles in other scientific practices such as software (Lamprecht et al. 2020; Hasselbring et al. 2020; Katz et al. 2021; Hong et al. 2022;Barker et al. 2022). While this framework is designed to promote transparency and data sharing, it remains subject to interpretation and implementation by individual data repositories, with potential variations in compliance and enforcement across different contexts (Boeckhout et al.2018; Tacconelli et al. 2022) and a prioritisation of machine readability over human inclusivity (Sterner and Elliot 2022).

GISAID’s approach to sound data management is also arguably compatible with FAIR principles but places a higher premium on regulating data flows and interactions between users and the infrastructure, as exemplified by its database access agreement. The EpiFlu™ Database Access Agreement is a mechanism designed to facilitate the sharing of influenza gene sequence data among researchers and public health professionals worldwide. This agreement outlines the terms under which users may provide data to the database, as well as the rights and obligations of authorised users with respect to that data. In particular, the agreement grants GISAID and authorised users a non-exclusive, worldwide, royalty-free, and irrevocable license to use, modify, display, and distribute the data submitted by users for research and intervention development purposes, provided they acknowledge the originating and submitting laboratories as the source of the data. Moreover, the agreement establishes certain restrictions on data access and distribution to ensure that users are acting in the best interests of public health. For example, users are not permitted to access or use the database in connection with any other database related to influenza gene sequences, nor are they allowed to distribute data to any third party other than authorized users. Users are also required to make best efforts to collaborate with representatives of the originating laboratory responsible for obtaining the specimen(s) and involve them in such analyses and further research using such data (GISAID, 2023). Although this agreement is established to promote collaboration between scientists, a recent publication in Science exposed GISAID as having different tiers of access which aren’t defined in the agreement (Enserink and Cohen 2023). ^2^

The differences in the interpretation and implementation of sound data management by GISAID and INDSC illustrate the pluralistic nature of data governance and highlight the need for critical reflection on the normative foundations and ethical implications of data sharing practice. Even though both data infrastructures share common epistemic goals, cognitive-cultural resources, and knowledge forms on complex biological systems, they create distinct digital artifacts as a result of different policies and values (Elliot 2022). A review of metadata by Bernasconi et al. (2021) conclude that, while GISAID’s partially closed model is likely to attract international collaboration from under-resourced countries, it fails to provide features of data provenance such as persistent URLs to samples or publications. The urgency to better understand the epistemic role of these infrastructures – and those to come after it - is underscored by the work of Chen et al. (2022) and Brito et al. (2022) who identified that countries in lower income groups often lack efficient genomic surveillance capabilities, not due to being able to access the data infrastructure but due to socioeconomic factors such as inadequate infrastructure, low national GDP, and meagre medical funding per capita.

Building on prior work on data sharing strategies and understandings of openness between these two systems of data governance (Leonelli 2021; Leonelli 2023), our aim in this paper is to provide an empirical examination of a crucial process of relevant data journeys (Leonelli and Tempini 2020), specifically the transfer of SARS-CoV-2 genetic data from the collection stage to the analytical stage. These data movements often cross institutional and international borders, thereby posing challenges to conventional scientific divisions of labour, disciplinary boundaries, and epistemic hierarchies. Despite the inherent challenges in identifying and reconstructing these journeys, they present valuable units of analysis for mapping and comparing the diverse practices and circumstances involved in the mobilization and utilization of data (Leonelli 2016a; Leonelli and Williamson 2023). At the heart of our inquiry lies the research question of how data infrastructures function as entities that mediate the interplay between data and research practices, thereby affecting the processes and outcomes of data exchange. To answer this, our methodology entails a synthesis of quantitative methods such as data collection, frequentist statistics and network analysis, and is informed by critical data studies debates on the governance and inclusivity of data infrastructures (Beaulieu et al. 2013; Kitchin 2014; Kitchin & McArdle 2016; Leonelli 2016b; Borgman 2017; Fecher 2018; Borgman and Bourne 2022; Wilson et al. 2022; Curry 2022).

## 3 Data and Methods

To explore collaborative research patterns across each data repository, our study draws on an analysis of collaboration in published articles and other bibliometric indicators. Bibliometrics as a meta-science methodology has been widely used to study the impact of COVID-19 on the research landscapes (Chahrour et al., 2020; Mohabab et al., 2020; Yinka Akintunde et al., 2021; Wang and Tian 2021; Acciai et al., 2022; Benach et al., 2022; Zhang et al., 2022; Zhong et al., 2022; Sofi-Mahmudi et al,. 2023). Although bibliometrics was established in the 1950’s, bibliometrics is increasing in usage across academic disciplines, and has become a central methodology to explore trends in international and national collaboration, thematic clustering of keywords and topics and structural patterns of networks dynamics (Subramanyam 1983; Donthu et al., 2021). By focusing on collaboration between institutions, countries and researchers, bibliometrics are able to better explore the involvement of actors involved in COVID-19 research. One caveat of this methodology is that it only captures a partial picture of research collaboration, with little information being placed on practices or informal collaborations, therefore it should not be understood as the entire landscape (Leonelli 2021). In recent years, a number of online databases such as Web Of Science (RRID:SCR_022706), Dimensions.ai (RRID:SCR_021977), have become easily accessible in providing a systematic collection of multi-disciplinary publications and a number of software packages such as VosViewer (RRID:SCR_023516), Bibliometrix (RRID:SCR_023744) and Gephi (RRID:SCR_004293) have made the analysis of such data more achievable (Muñoz et al., 2020).

Each corpus was analysed using common bibliometric indicators, collaboration and equity measurements and network analysis. To begin, we considered indicators which are grounded in the existing bibliometric literature such as total number of publications, average citations per publication, total citations, average Altmetric score, accessibility options (Open or Closed Access), publishers’ landscape, and co-occurrence of key terms within publications. The results of these were plotted as line graphs or tree maps for each dataset. After this, we extended our statistical framework to include a bar chart of the distribution of variants between each dataset - aligning our approach with many bibliometric studies that focus on phenomena-specific metrics. We then deployed two measurements to understand the author and income collaborations between countries. The first is based on inter and intra-regional collaborations and the second identifies collaboration in relation to income groups classified by the World Bank.

Lastly, we explored the relational dynamics among publications by engaging in bibliographic coupling and social network analysis (SNA). We employed the standard formulation of bibliographic coupling as introduced by Kessler in 1963:

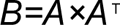

Here, B is the bibliographic coupling matrix, and A is the bipartite network adjacency matrix. Each row and column in A correspond to a node, which may represent entities such as publication data, countries, academic institutions, and funders. The elements a^ij^ in the adjacency matrix A represent the number of bibliographic couplings between article *i* and article *j*. We used two common SNA metrics to plot the structural impacts of the network: *degree and betweenness*, where degree is the number of associations a particular node has, and *betweenness* represents the number of occurrences a node acts as a bridge between one node to another. (Newman 2005). All analysis was conducted using an R Project for Statistical Computing (RRID:SCR_001905). The project, along with the code and data to reproduce the analysis have been made openly available on Zenodo (10.5281/zenodo.8399189)

### 3.1 Data Collection and Filtering

In bibliometric analyses, delineating precise search strategies and choosing a source of truth database is of paramount importance to capture a comprehensive dataset.

Within the genomic data repositories landscape, GISAID and the INSDC groups exhibit divergent data citation protocols. Crucially, GISAID enforces a stringent requirement whereby members must cite and formally solicit permission from data depositors prior to any publication endeavours. This procedural characteristic facilitates a streamlined search strategy for GISAID data. Specifically, by employing the terms “Global initiative on sharing all influenza data”, “GISAID” or “EpiCov” in conjunction with a curated set of COVID-19-centric keywords. Conversely, the INSDC groups, devoid of such stipulations, present a slightly more intricate search matrix. Though there is an absence of a formal citation requirement, it is a common observation that published outputs frequently reference the PRJ accession code associated with a sequence or directly name the data repository. Consequently, the search query for the INSDC groups necessitates the integration of terms such as “ENA”, “The Covid-19 Data Portal”, “NCBI Virus”, and “NiH” with additional accession codes such as “PRJ”. A full table of search queries can be found in table 1. In order to ensure the data queried was relevant to the study of SARS-CoV-2 and not any other influenza or pathogen disease, the following inclusion criteria are applied: A Covid-19 related term had to be in the full text; Study had to be an academic article; Published between January 2020 and January 2023; and all types of methods and disciplines are allowed.

**Table 1.** Workflow for Dimensions.ai Searches on Publications Referencing Major SARS-CoV-2 Data Repositories.

| Repository | Search Query | Covid-19 Related Terms | Additional Filters |
| --- | --- | --- | --- |
| GISAID | "GISAID" OR<br>"EpiCoV" OR<br>"Global initiative<br>on sharing all<br>influenza data" | • *"sars-cov-2"*<br>• *"covid"*<br>• *"covid-19"*<br>• *"coronavirus"*<br>• *"HCov-19"* | • Publication type must be a scientific article.<br>• Articles between January 1<sup>st</sup> 2020 and January 1<sup>st</sup> 2023.<br>• Articles can be from any discipline or methodology.<br>• Remove duplicates of DOI's.<br>• Remove articles where country affiliation is not documented. |
| NCBI | "NCBI Virus" OR<br>"genbank" OR<br>"National Center<br>for Biotechnology<br>Information" OR<br>"Accession: PRJ"<br>OR "International<br>Nucleotide<br>Sequence<br>Database<br>Collaboration" |  |  |
| ENA | "The Covid-19<br>Data Portal" OR<br>"European<br>Nucleotide<br>Archive" OR<br>"ENA" OR<br>"Accession: PRJ"<br>OR "International<br>Nucleotide<br>Sequence<br>Database<br>Collaboration" |  |  |
| DDBJ | "DDBJ" OR "DNA<br>Data Bank of<br>Japan" OR<br>"Accession: PRJ"<br>OR "International<br>Nucleotide<br>Sequence<br>Database<br>Collaboration" |  |  |

Using these search queries and filters, we choose to use the Dimensions Analytics API (Hook et al., 2018; Herzog et al., 2020) as our *source of truth* to build collections for each repository. Established in 2018 by Digital Science, the Dimensions Analytics API provides one of the largest sources of publication data (Adams et al., 2018; Bode et al., 2019; Visser et al., 2020). Our initial search returned the following number of publications for each repository: GISAD (*n = 14092), NCBI (n = 12751)*, *ENA (n = 13491)*, *DDBJ (n = 508)*. However, as Guerrero-Bote et al., 2021 point out, the validity of Dimensions data can often be less reliable to Scopus – as it depends on machine learning curated data and fields such as research affiliations often contain missing entries and may contain duplicates based on preprints. For these reasons, we further filter our data to remove any duplicates of publication.ids or titles and for our network analysis we remove any entries where country or institution affiliation is not documented. After this filtering, the size of our final dataset for each repository is: GISAD (*n = 11945), NCBI (n = 10685)*, *ENA (n = 9728)*, *DDBJ (n = 325)*. Similarity between each corpus’s publication id – shown in figure 1 – identifies that each repository has a mostly unique corpus of data. The biggest overlap between corpuses was between GISAID and NCBI, narrowly followed by the ENA and NCBI. There was a significantly low similarity of articles mentioning all three INSDC members.

**Figure 1.**
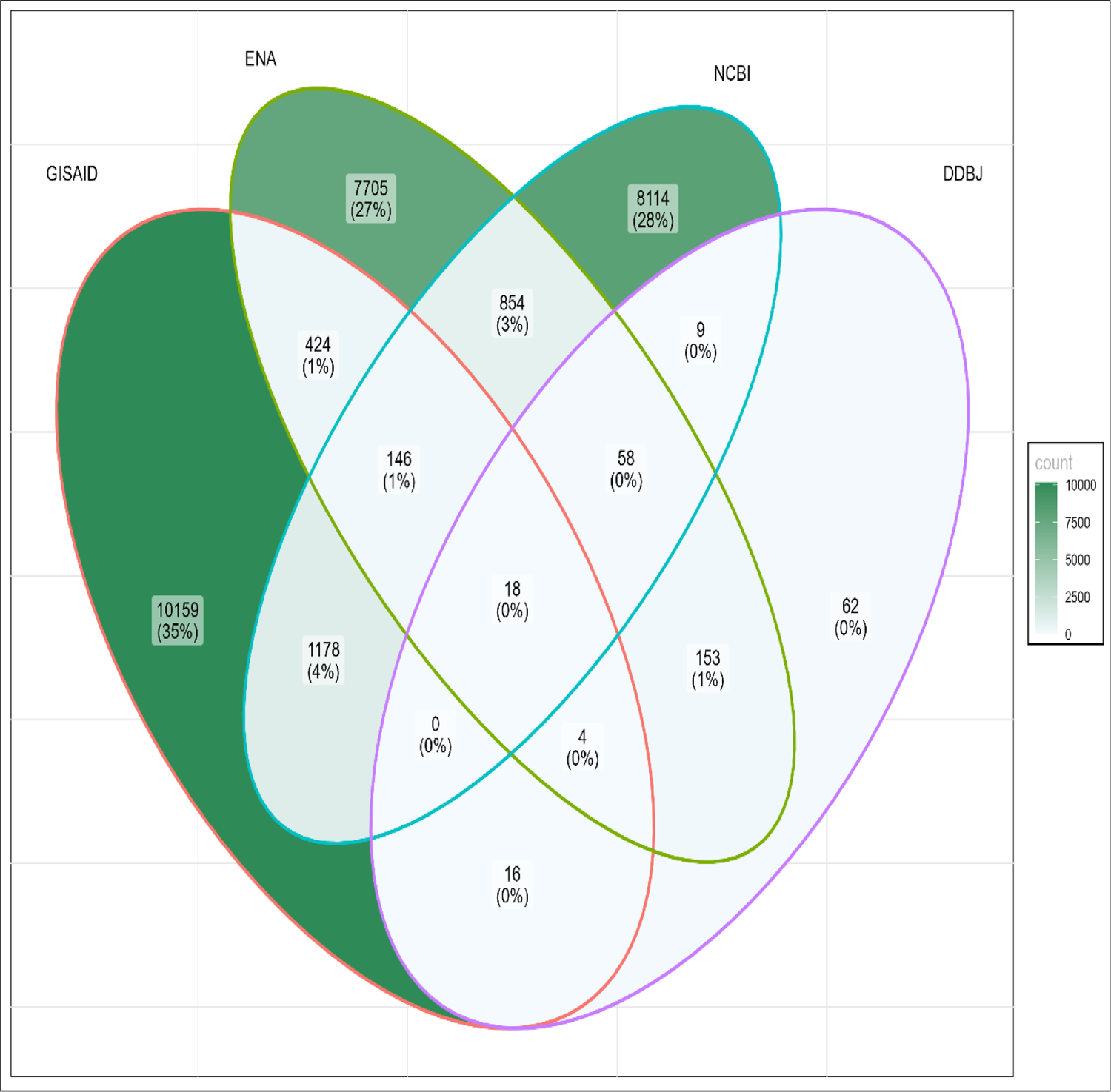
Corpus Similarity Across Major SARS-CoV-2 Repositories: Similarity of Publication Corpus’s Across Major SARS-CoV-2 repositories (January 2020-January 2023). Each oval represents a repositories corpus with GISAID (red), ENA (green), NCBI (blue) and DDJB (purple). The total number of unique publications are labelled for each overlapping circle as well as the percentage of the entire corpus.

## 4 Results

### 4.1 Bibliometric Indicators

Notwithstanding the comparatively reduced aggregate of publications in the GISAID repository relative to the comprehensive corpus of the INSDC, GISAID manifests a superior metric in several key bibliometric indicators such as average citations per month, total monthly citations, and average Altmetric scores as seen in Figure 2.

**Figure 2:**
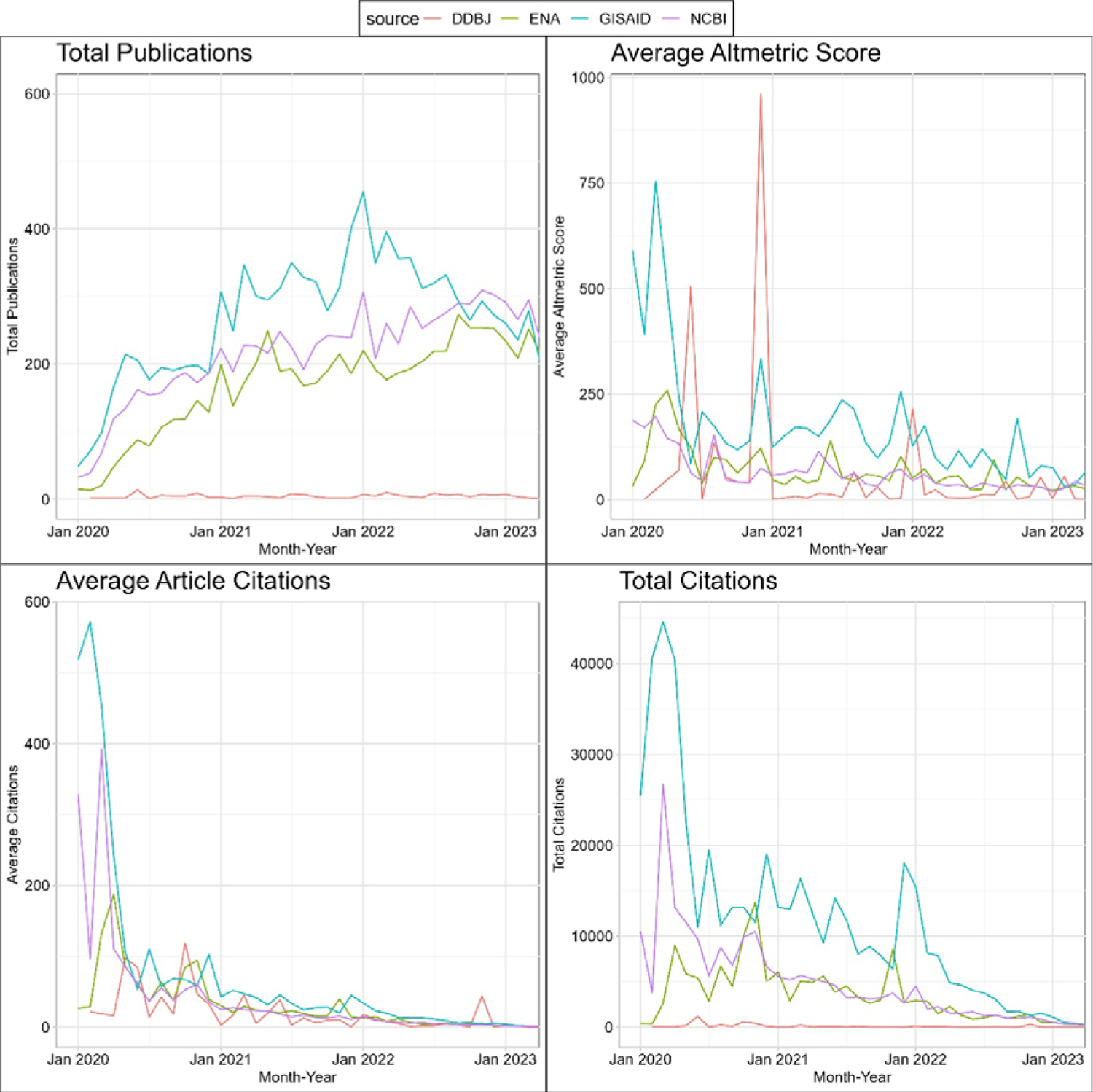
Temporal Trends in Scholarly Metrics Across Major SARS-CoV-2 Repositories. This line graph illustrates longitudinal variations in key bibliometric indicators: (1) Total Number of Publications (top left), (2) Average Citations per Publication (top right), (3) Average Altmetric Score (bottom left), and (4) Cumulative Citation Count (bottom right). Data points span from January 2020 to January 2023. The DNA Data Bank of Japan (DDBJ), National Center for Biotechnology Information (NCBI), European Nucleotide Archive (ENA), and Global Initiative on Sharing All Influenza Data (GISAID) are represented by red, purple, green, and aqua blue lines, respectively.

This discrepancy is largely attributable to GISAID’s recurrent citation in seminal works delineating the initial virological properties of SARS-CoV-2 (Coronaviridae Study Group of the International Committee on Taxonomy of Viruses, 2020; Corman et al., 2020; Zhu et al., 2020; Wang et al., 2020; Wrapp et al., 2020; Wall et al., 2020), discussions of its phylogenetic origins (Holshue et al., 2020; Zhou et al., 2020; Andersen et al., 2020), as well as foundational studies on therapeutic and vaccine protocols (Hoffmann et al., 2020; Polack et al., 2020; Wölfel et al., 2020).

In the context of the INSDC consortium, the total number of publications across all databases stands at *n=20,059*, which is fifty nine percent more than GISAID with *n=11,945*. Among the INSDC constituents, DDBJ records the most modest performance across all bibliometric impact measures. However, the DDBJ makes a perk in the Altmetric graph, this spike reflects both the low number of publications mentioning the DDBJ but also the high alt metric score of Amendola et al., 2020’s early work on evidence of SARS-CoV-2 RNA from a swab in Italy December 2019. Conversely, despite possessing a fewer number of articles than NCBI, the ENA surpasses all other INSDC members impact factors, facilitated in part by pivotal contributions to the PRIDE database resource (Perez-Riverol et al., 2022) and the Ensembl 2021 project (Howe et al., 2020).

In Figure 3 for all GISAID, NCBI, ENA and DDBJ repositories, Gold Open Access (OA) emerges as the predominant access modality, registering a prevalence of 43%, 47%, 39% and 71% respectively. In GISAID, subsequent access modalities adhere to a conventional OA hierarchy, featuring Green (32%), Bronze (12%), Hybrid (8%), and Closed (5%) categories. ENA diverges the most, with Closed (25%) being the second leading access type, followed by Green (18%), Hybrid (11%) and lastly Bronze (8%). For the NCBI access types, Green (20%), Closed (16%), Bronze (8%) and Hybrid (8%) trail Gold. The case of DDBJ is a similar pattern, with Green (20%), Closed (15%), Bronze (5%) and Hybrid (8%).

**Figure 3.**
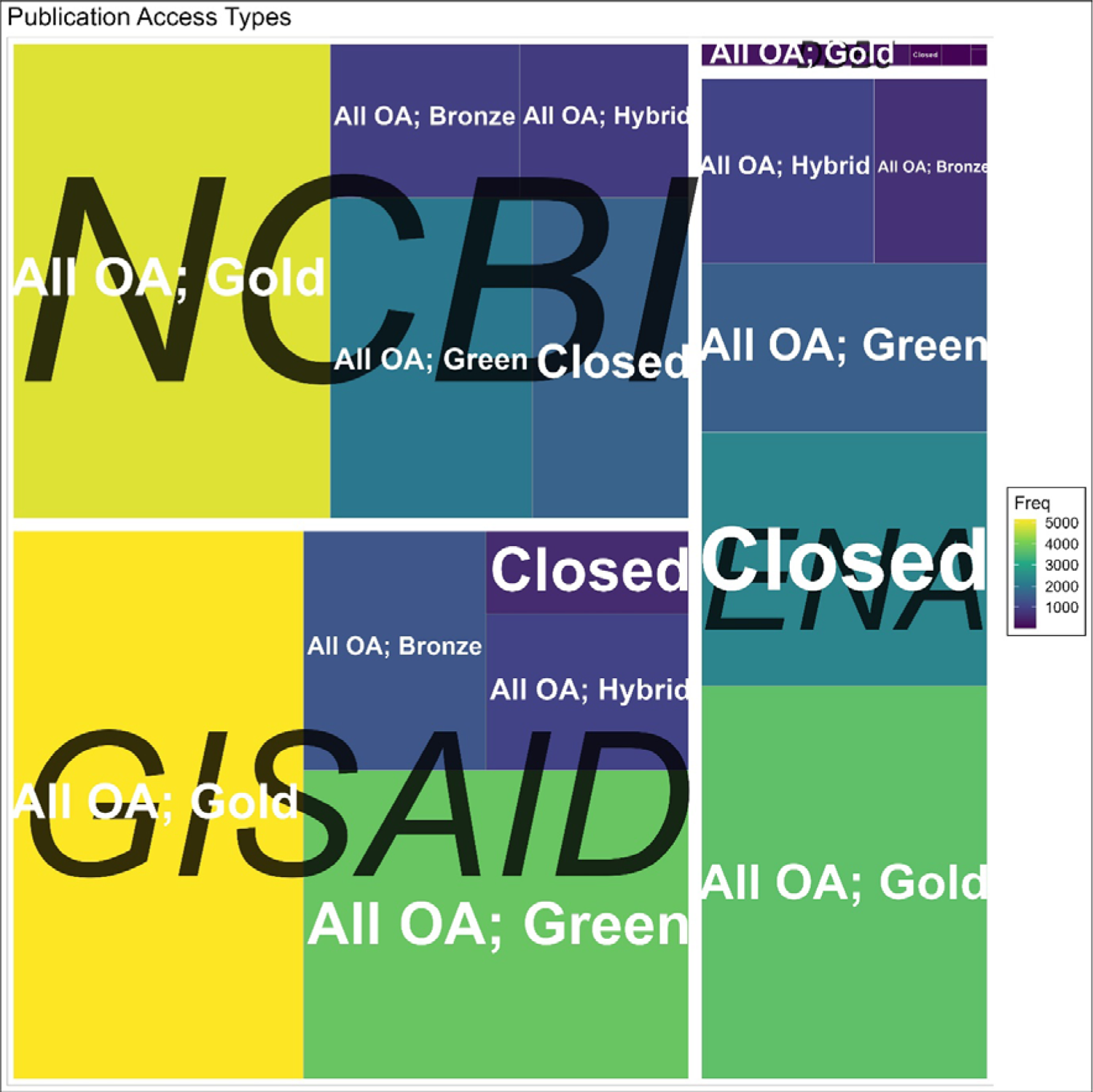
Publication Access Types Across Major SARS-CoV-2 Repositories: This treemap quantifies the distribution of open access (OA) – Gold, Green, Bronze and Hybrid - and closed access publications across The DNA Data Bank of Japan (DDBJ), National Center for Biotechnology Information (NCBI), European Nucleotide Archive (ENA), and Global Initiative on Sharing All Influenza Data (GISAID). Each rectangle’s area is proportional to the frequency of publications within that category. Databases are delineated by white borders and labelled at their centres in italicized black text. Within each database, access type are labelled in white text.

For NCBI in Figure 4, publishers in terms of prominence are Elsevier (19%), Springer Nature (16%), MDPI (12%), Cold Spring Harbor Laboratory (11%), Frontiers (8%). In the same figure, ENA exhibits a similar trend with Springer Nature (27%), Elsevier (20%), MDPI (8%), Cold Spring Harbor Laboratory (8%), and Frontiers (5%). In the case of DDBJ, the leading publisher instead is Centre for Disease Control and Prevention (41%), then followed by Springer Nature (14%), Elsevier (9%), Oxford University Press (6%) and MDPI (6%). GISAID demonstrates a distinct preference for Elsevier (20%) trailed by Cold Spring Harbor Laboratory (20%), Springer Nature (12%), MDPI (9%), and Research Square Platform LLC (5%). All other publishers were less than five percent.

**Figure 4.**
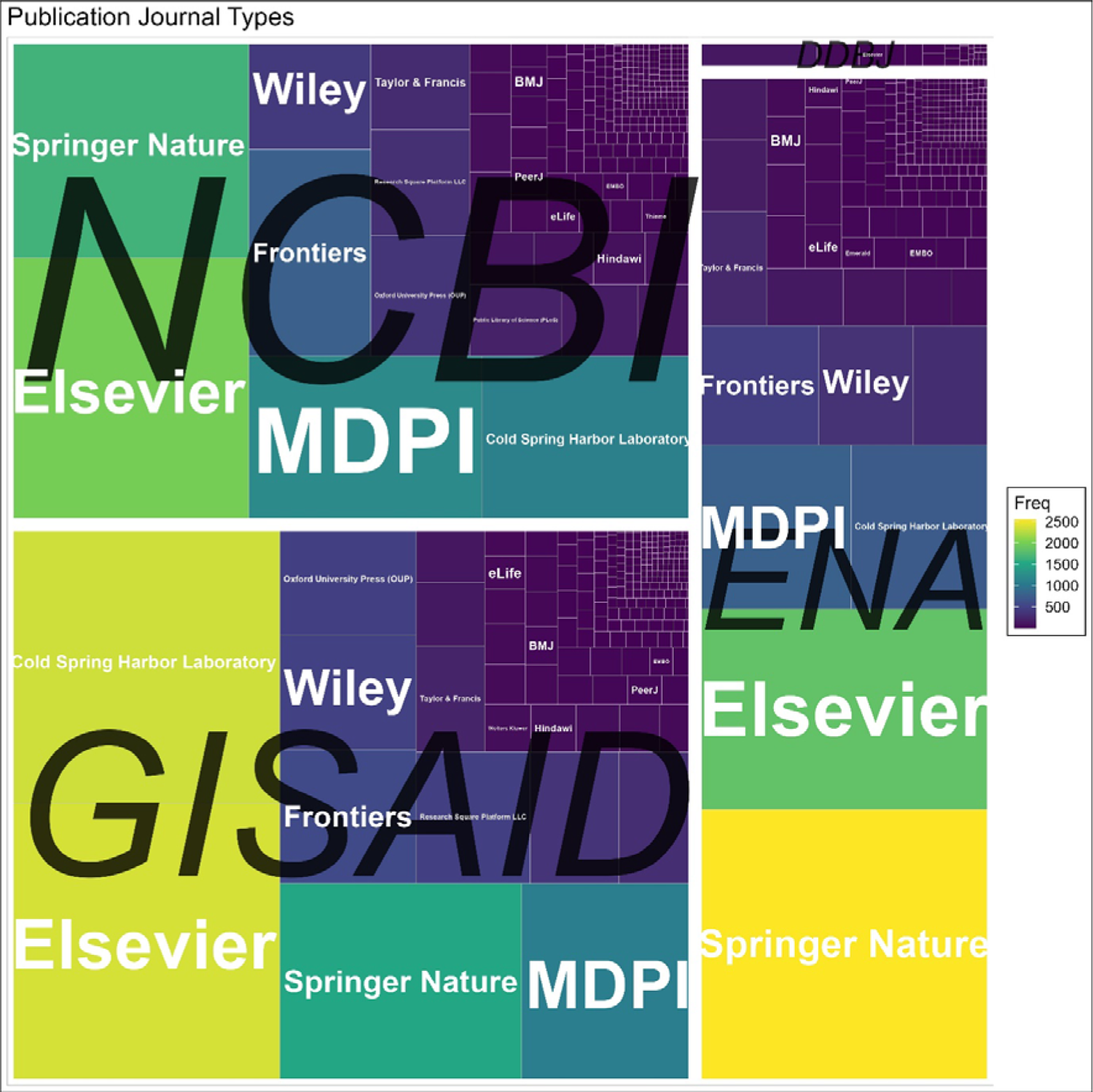
Publisher Landscape Across Major SARS-CoV-2 Repositories. This treemap quantifies the distribution of publishers across The DNA Data Bank of Japan (DDBJ), National Center for Biotechnology Information (NCBI), European Nucleotide Archive (ENA), and Global Initiative on Sharing All Influenza Data (GISAID). Each rectangle’s area is proportional to the frequency of publications within that category. Databases are delineated by white borders and labelled at their centres in italicized black text. Within each database, publishers are labelled in white text.

### 4.2 Variant and Keyword Distribution

For all repositories, the World Health Organization (WHO) nomenclature exhibited a more pervasive mention as opposed to the Pango lineage classifications as shown in figure 5. In the ENA corpus, variant denominations predominantly adhere to the WHO taxonomy, with the following distribution: Alpha (145), Beta (145), Delta (142), Gamma (54), and Kappa (20). Conversely, the Pango lineage descriptors manifest with a diminished frequency: p.2 (75), B.1.1.7 (49), B.1.617.2 (32), B.1.351 (23), and B.1.1.529 (21). GISAID encapsulates the largest share of variant mentions. Predominantly, WHO nomenclature appear more regularly, with leading variants being Omicron (2093), Delta (1879), Alpha (1098), Beta (795), and Gamma (606). For GISAID, Pango lineages appear with B.1.1.7 (832), B.1.351 (506), B.1.617.2 (514), P.1 (400), B.1.1.529 (367). The NCBI encompasses the most substantial proportion of variant mentions among the INSDC repositories. The paramount WHO classifications are Alpha, Beta, Delta, Omicron, and Gamma with frequencies of 274, 263, 252, 239, and 136 respectively, and for Pango lineages, they are B.1.1.7 (86), P.1 (77), B.1.351 (65), B.1.617.2 (60), and B.1.1.529 (34).

**Figure 5:**
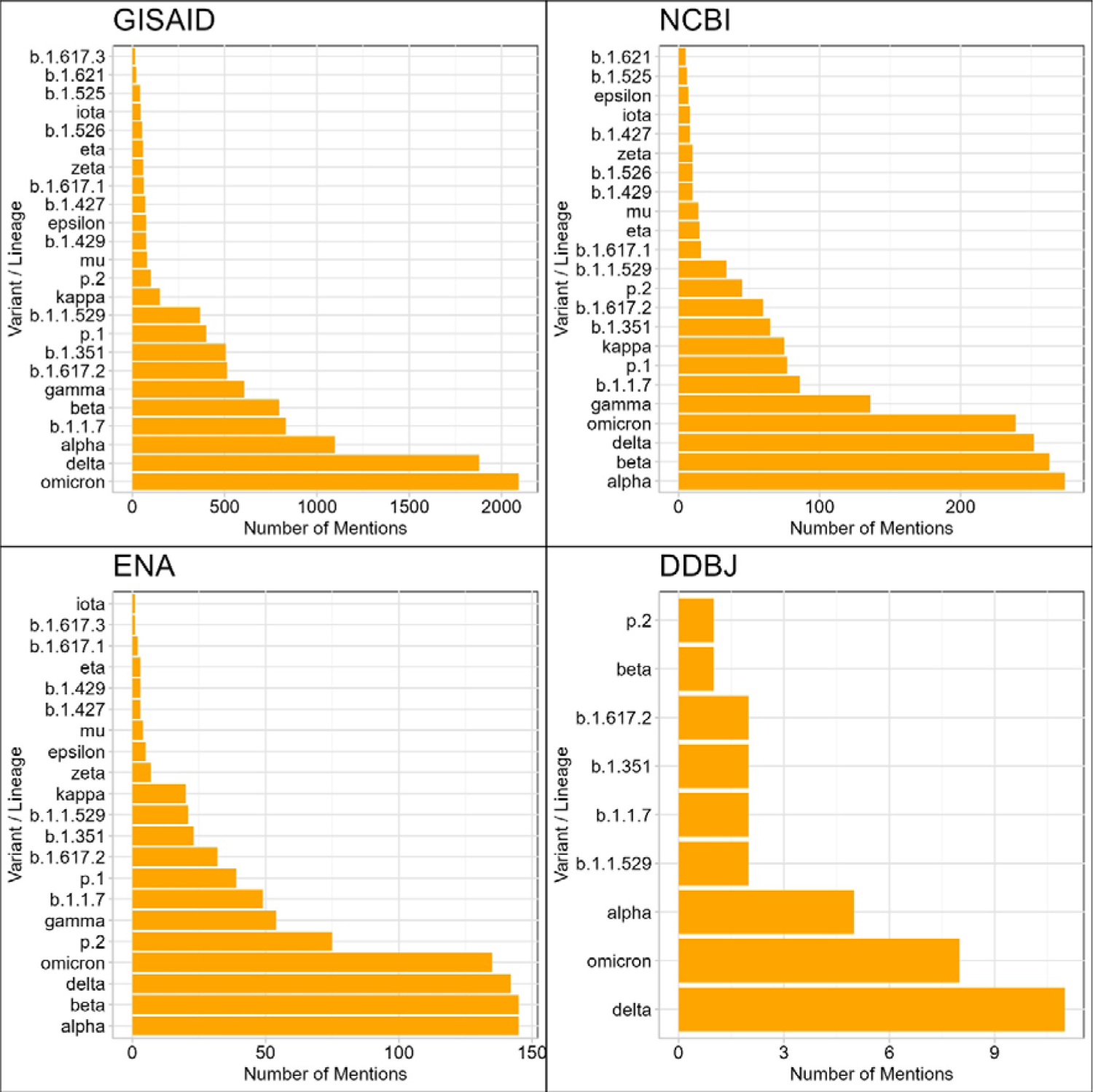
Frequency Distribution of Viral Variants Mentioned in Abstracts Across Major SARS-CoV-2 Repositories. This bar chart represents the frequency of mentions for specific viral variants and lineages in the dataset’s abstracts. WHO labels (e.g., alpha, beta) and Pango lineages (e.g., b.1.1.7, b.1.351) are accounted for. Each bar corresponds to a distinct variant or lineage, ordered in descending frequency of mentions. The x-axis quantifies the number of abstracts mentioning each variant, and the y-axis identifies the respective variants and lineages.

Figure 6 shows that the keyword distribution of Medical Subject Headings (MeSH) terms convergence on analogous themes across the repositories, encompassing gender dichotomies (female and male), species delineations (animals, human, mice), age categorizations (adult), data types (RNA, genomics, genome), and facets of pandemics and viral evolution (pandemics, viral, spike glycoprotein, phylogeny). An idiosyncratic characteristic of GISAID is its heightened frequency of therapeutic-centric terminologies (vaccines, mutations, neutralising antibodies). In juxtaposition, other INSDC repositories underscore methodological lexemes such as databases and computational biology. Subsidiaries of INDSCINSDC spotlight repository-specific keywords: for instance, the ENA emphasises Climate Change, while the DNA Data Bank of Japan (DDBJ) accentuates infections, pneumonia, and, not surprisingly, Japan. Within the confines of GISAID and NCBI, the uppermost fifteen keywords converge on a centrality of fourteen. In contrast, DDBJ and ENA oscillate between a centrality range of five to twelve and a half. Within the network framework of GISAID, the highest betweenness centrality is attributed to “adult” and “neutralizing.” Conversely, for NCBI, “computational biology” and “mice” emerge preeminent, for ENA it’s “databases” and “male”, and for DDBJ, “humans” and “animals” occupy this distinction.

**Figure 6:**
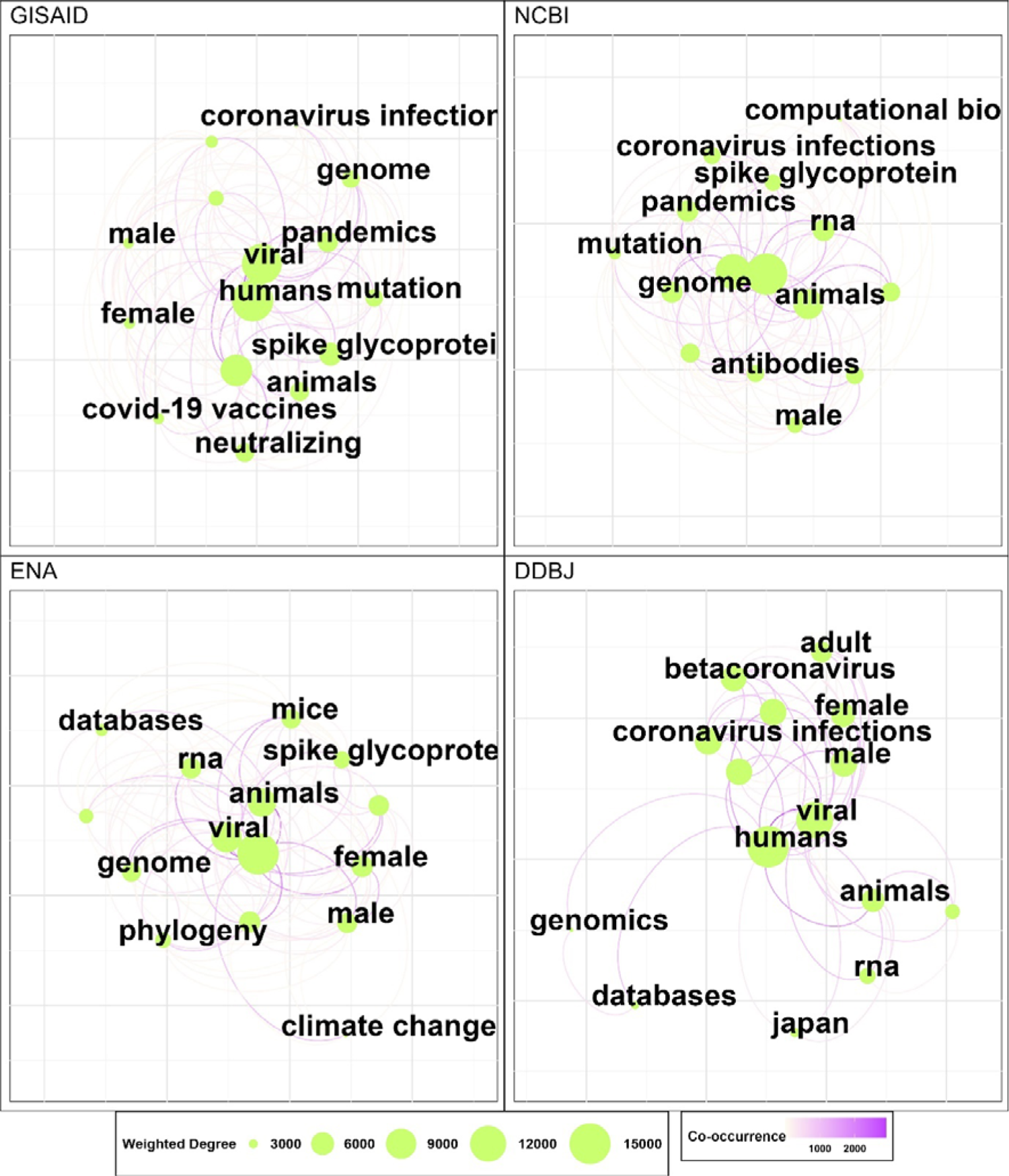
Co-Occurrence Network of Top MeSH Terms Across Major SARS-CoV-2 Repositories. This graph depicts a co-occurrence network of the top 15 *Medical Subject Headings (MeSH) terms within the given dataset, represented as nodes. Edges between nodes are weighted by the frequency of co-occurring terms across publications, and edge thickness scales with weight; edges with weight below a threshold of 5 are excluded for clarity. Node size is dictated by the node’s degree, and colour is mapped to betweenness centrality*.

### 4.3 Author Collaboration

As table 3 shows, within the GISAID repository, the year 2020 manifested a marked inclination towards collaborative endeavours, evidenced by a significant number of papers with either 2-5, 5-10, and 10-20 authors juxtaposed against a modest 64 single-authored papers. By 2021, a proliferation in contributions was observed, most prominently in the 10-20 author category, amassing a noteworthy 1,207 papers.

However, by 2022, while the papers in the 10-20 author bracket peaks at 1,254, those exceeding 50 authors witnessed a slight decrement to 60. In terms of hyper authorship, GISAID featured a decent share of 50-100 authors and was the leading year-on-year growth of 100-250 authors. Although GISAID didn’t have any papers with between 250-500 authors, they beat every member of the INSDC in the extreme authorship categories with leading representation every year for 500-1000 and 1000-5000.

The ENA database, in 2020, registered most of its papers within the 2-5 author cohort with 420 contributions. This was contrasted by 72 papers in the single author category and 29 in the 50+. An intriguing surge was witnessed in 2021, particularly with a staggering 1,219 in the 2-5 authored contributions and related increases in all other categories. In 2022, the surge continued but at a slower rate, with the most significant increase being the jump in single author papers to 407. The ENA trumped GISAID in the 50-100 author per paper and had the best year on year representation for the hyper authorship categories between the INSDC members. For the NCBI repository, the data from 2020 underscores a dominant trend towards collaborative work, with 613 papers in the 2-5 author bracket and 546 in the 5-10 author segment. This collaborative proclivity augmented in 2021, especially within the 2-5 author category which culminated in 1,032 papers. In 2022, all categories up to 50 authors continued to rise. NCBI displayed marginally less than the ENA in the 50-100 authorship category and marginally less than GISAID in the 100-250 authorship category but failed to represent continuous improvement in hyper authorship categories on a yearly basis. Lastly, the DDBJ repository, in 2020, was most prolific in the 2-5 author segment, recording 49 papers. The subsequent year, 2021, saw relatively modest numbers, with the 2-5 author category still leading but with less papers. However, 2022 registered a marginal upswing, particularly in the 2-5 and 10-20 author category. By 2023, the repository maintained a only had one paper with a hype authorship of 50+ across all years.

### 4.4 Geographical and Income Collaboration

Figure 7 shows, the ENA was the only repository to have a higher share of multi-region collaborations than single region, with 13,819 beating 15,269. Countries with the leading number of documents were United States, United Kingdom, Germany, China, Australia, Spain, Italy, India, France, Canada, Netherlands, Japan, Switzerland, Brazil, Sweden, South Africa, Belgium, Austria, Denmark, and Norway. Countries with a higher ratio of multi-region collaborations were South Africa, Canada, Australia, Sweden, Brazil, United Kingdom, Belgium, Denmark, Japan, United States, Norway, Spain, Austria, and the Netherlands. Countries in the ENA with a large proportion of single region collaborations were China, Italy, France, and India. Germany had a relatively equal ratio between the two collaborations.

**Figure 7:**
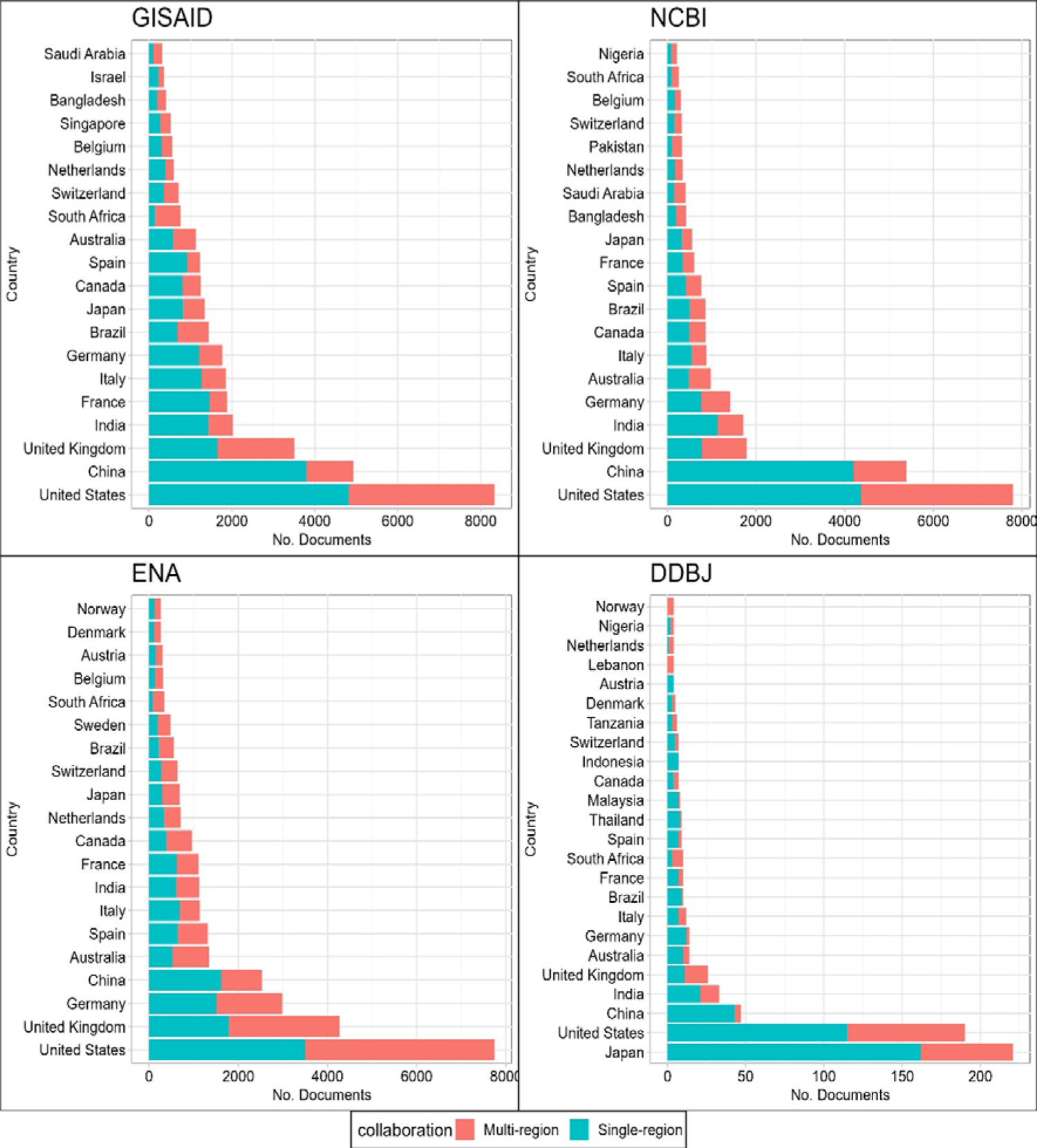
Distribution of Single- and Multi-Region Collaborations in Scholarly Publications by Top 20 Countries Across Major SARS-CoV-2 repositories. *This stacked bar chart portrays the extent of single-region and multi-region collaborations in scholarly documents for the top 20 countries based on publication volume. The x-axis indicates the number of documents associated with each country, and the y-axis lists the countries in descending order of total documents. The colours in each bar segment represent the type of collaboration: single-region or multi-region*.

**Figure 8:**
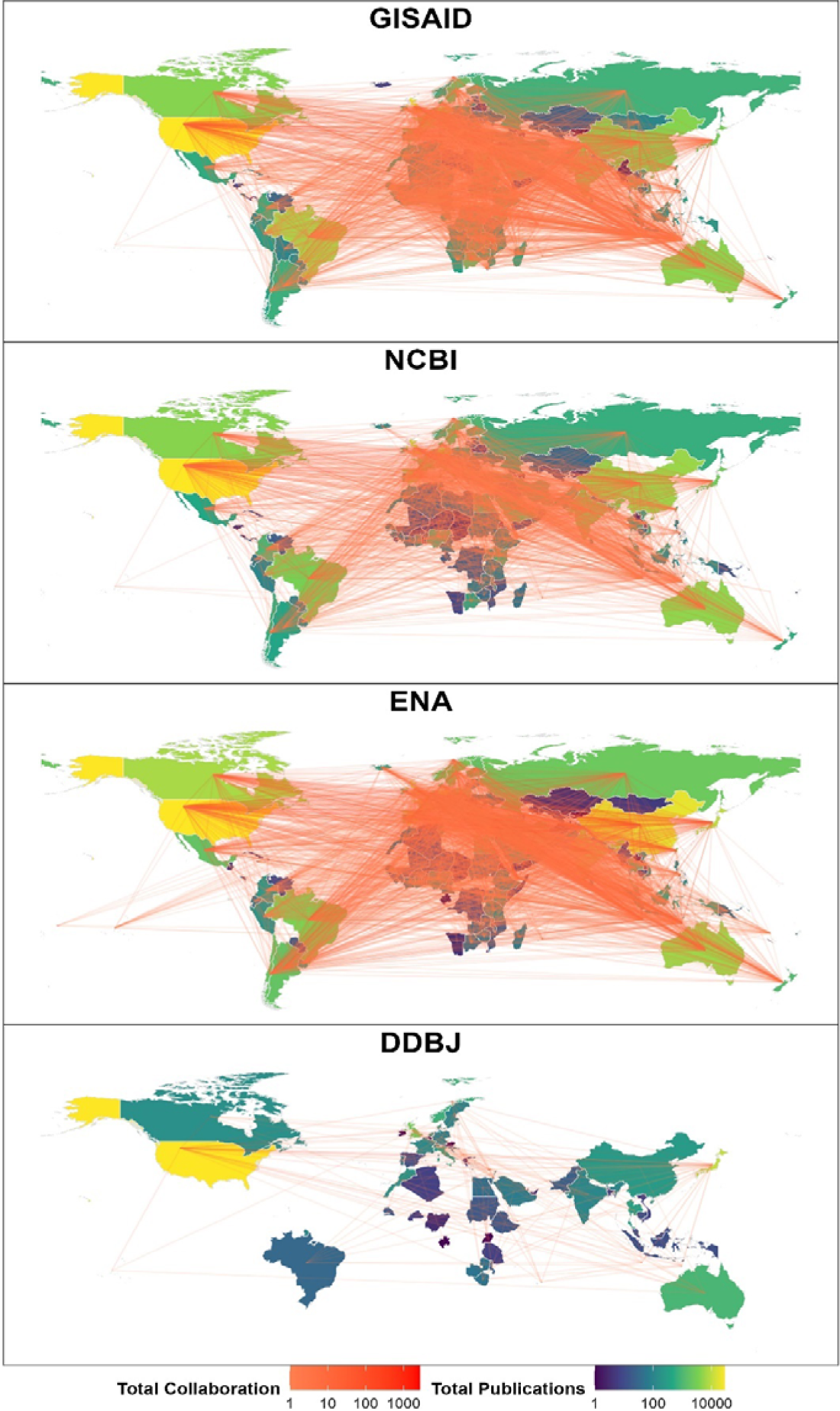
Country Collaborations Across Major SARS-CoV-2 Repositories. The figure provides a visual representation of global collaborations among countries. It plots a geographical map overlaid with collaboration lines between countries, where the colour of the lines represents the intensity or weight of the collaborations.

**Figure 9:**
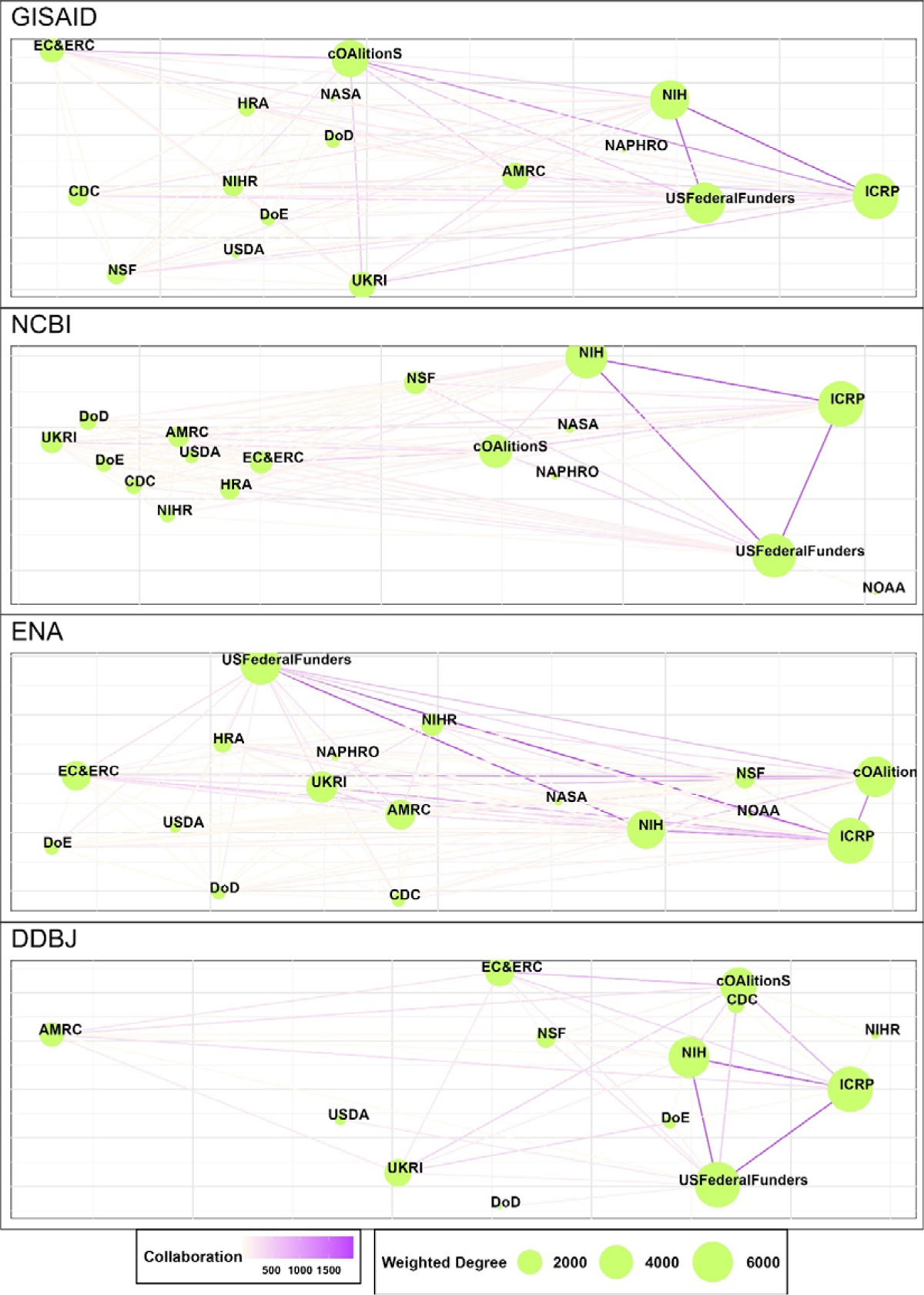
Funder Group Collaborations Across Major SARS-CoV-2 Repositories: *This graph depicts a collaboration network of funder groups within the given dataset, represented as nodes. Edges between nodes are weighted by the frequency of co-occurring funders across publications. Node size is dictated by the node’s weighted degree*.

**Figure 10:**
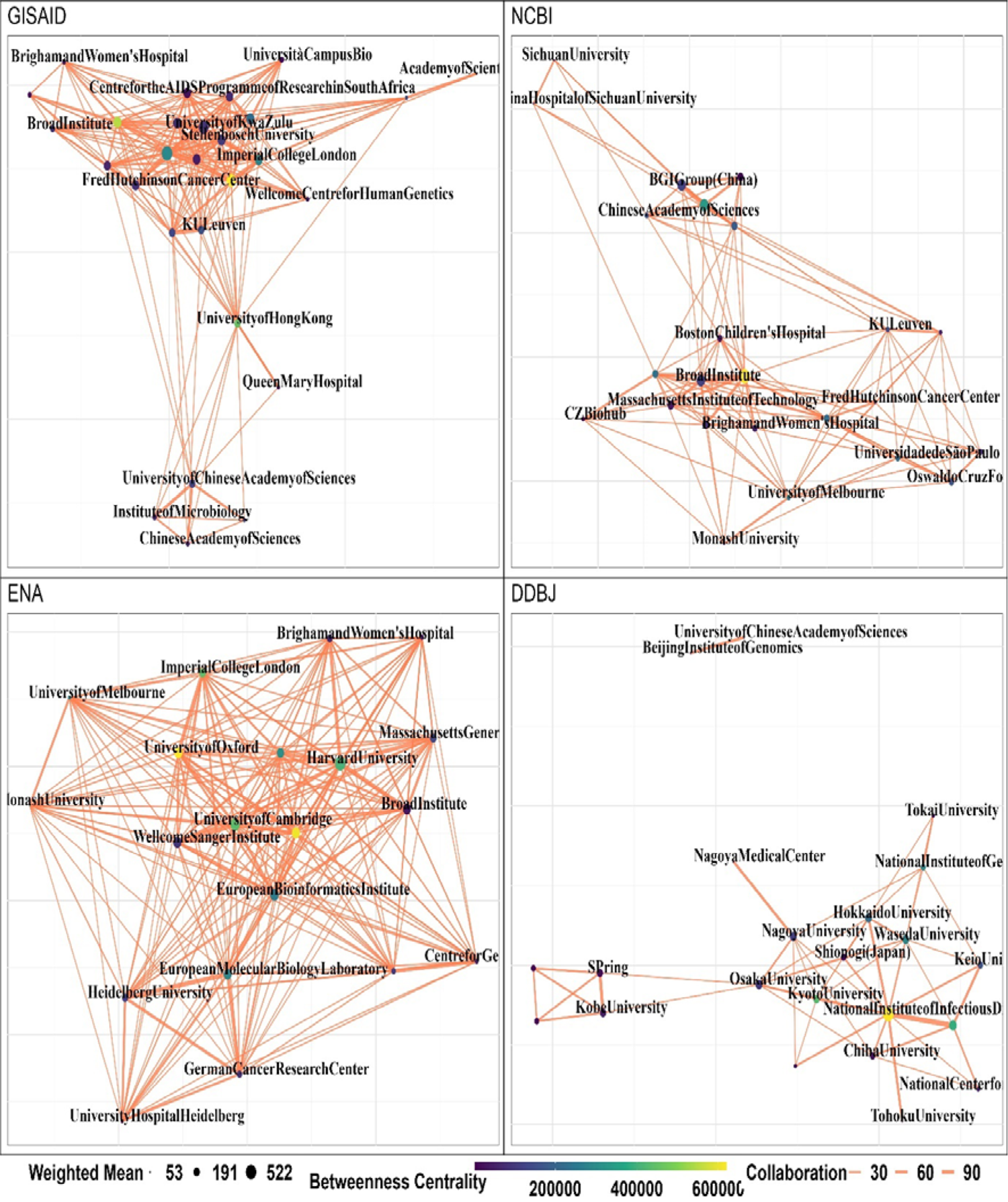
Institution Group Network Across Major SARS-CoV-2 Repositories. This graph depicts a collaboration network of the institutions within the given dataset, represented as nodes. Edges between nodes are weighted by the frequency of collaborating institutions across publications. Node size is dictated by the node’s weighted degree, and colour is mapped to betweenness centrality, following a viridis scale.

For GISAID single region collaborations were the most frequent, with a total of 21,432 over 13,476 multi-region collaborations with the total number of documented coming from United States, China, United Kingdom, India, France, Italy, Germany, Brazil, Japan, Canada, Spain Australia, South Africa, Switzerland, Netherlands, Belgium, Singapore, Bangladesh, Israel, and Saudi Arabia. The countries with the most amount of multi-regional collaboration were South Africa, United Kingdom, Brazil, and Bangladesh, all of which had more multi-regional collaborations than single-region. Following this, Switzerland, Singapore, and Australia had roughly equal share between the two collaborations categories. Countries with the highest single region collaboration were France, China, Spain, India, Italy, and Germany, all of which had over double the amount of single regional collaborations over multi-regional collaborations. Netherlands, Saudi Arabia, Canada, Japan, Israel, and United States had between 40-10% more single regional collaborations.

For NCBI the split between single region and multi-region collaboration were less than GISAID, with 15,440 and 10,742 respectively. The leading countries with the greatest number of documents were United States, China, United Kingdom, India, France, Italy, Germany, Brazil, Japan, Canada, Spain, Australia, South Africa, Switzerland, Netherlands, Belgium, Singapore, Bangladesh, Israel, and Saudi Arabia. Countries with the highest multi-regional collaboration were Brazil, South Africa, Saudi Arabia, Australia, Bangladesh, and United Kingdom. Countries with the highest single-regional collaborations were China, India, Italy, Germany, Canada, Japan, and France. The rest of the countries had a relatively equal ratio of single and multi-region collaborations.

DDBJ had the lowest number of total articles and had greater share of single region, 451, collaborations over multi region 214. The leading countries collaborating on documents were Japan, United States, China, India, United Kingdom, Australia, Germany, Italy, Brazil, France, South Africa, Spain, Thailand, Malaysia, Canada, Indonesia, Switzerland, Tanzania, Denmark, Austria, Lebanon, Netherlands, Nigeria, and Norway. Lebanon had the entire of its collaborations based as multi-region, with the Netherlands, South Africa and the United Kingdom following them with higher proportions of multi-region collaboration. China, Brazil, Thailand, Malaysia, Germany, Spain, Japan, Australia, Switzerland, France, India, United States, Denmark, and Canada all had over 40% more single regional collaborations, with the rest of the countries having a tied split.

From the data presented in Table 2, it is evident that collaborations between High-Income-to-High-Income groups (HI-HI) dominate the submissions in GISAID, NCBI, and ENA with notable figures of 8504, 6599, and 11199 respectively. DDBJ shows a significantly lower count of 151 in this category. In contrast, collaborations between Low Middle Income and Low Income (LMI-LI) groups are sparse across all repositories. Interestingly, collaborations involving Upper Middle-Income groups (UMI) with other income groups, such as Low-Income (LI) and Low Middle Income (LMI), show varied results across databases. For instance, UMI-LMI collaborations are relatively higher in NCBI (349) compared to GISAID (269) and ENA (228), while DDBJ shows little representation in this category. Furthermore, an intriguing pattern observed in the other HI category, where each repository – apart from the DDBJ – leads the collaboration type in one way, most interestingly the ENA database exhibits a substantially higher number of collaborations in the HI-MIX category (3830) compared to both GISAID (2099), NCBI (1754), GISAID led in the HI-UMI category (6494) followed by the ENA (5376) and the NCBI (4521). Both NCBI (2214) and GISAID had the same number of HI-LMI of 2161, followed by the ENA 1724. The DDBJ does feature as the greatest percentage 21.7% but with significantly less collaborations.

**Table 2.** Income Collaboration Across Major SARS-CoV-2 Repositories. This table presents the share of income group (based on World Bank classifications) collaborations for each repository.

| **Income Collaboration** | *GISAID* | *NCBI* | *ENA* | *DDBJ* |
| --- | --- | --- | --- | --- |
| *HI-HI* | 7894 (41.3%) | 6340 (41.5%) | 10372 (47.7%) | 151 (64.3%) |
| *HI-LI* | 138 (0.72%) | 113 (0.74%) | 163 (0.8%) | 6 (2.6%) |
| *HI-LMI* | 2161 (11.3%) | 2161 (14%) | 1724 (8%) | 51 (21.7%) |
| *HI-MIX* | 2099 (11%) | 1754 (11.2%) | 3830 (24.7%) | 14 (6%) |
| *HI-UMI* | 6494 (34%) | 4521 (30.3%) | 5376 (26%) | 49 (20.9%) |
| *LMI-LI* | 4 (0.02%) | 19 (0.15%) | 21 (1%) | 2 (0.9%) |
| *UMI-LI* | 17 (0.09%) | 16 (0.1%) | 13 (0.06%) | 12 (5.1%) |
| *UMI-LMI* | 269 (1.41%) | 349 (2.28%) | 224 (1%) | 0 (0%) |
| *UMI-MIX* | 25 (0.13%) | 14 (0.1%) | 19 (0.09%) | 0 (0%) |

### 4.5 Networks

Table 3 provides a comprehensive network analysis of the various databases, detailing their network characteristics across different types: Authors, Country, Funder, and Institution. A salient observation from the table is the pronounced homogeneity in the funding network across the databases. Specifically, the nodes in this network span from 13 (DDBJ) to 17 (both ENA and NCBI), with GISAID closely trailing at 16. GISAID’s funding ecosystem, with a mean weighted degree of 2289, indicates a potential proclivity of funders towards research utilizing GISAID data. Nonetheless, the elevated clustering coefficient and density metrics for all databases underscore a robust collaboration among funders.

**Table 3.** Comparative Network Statistics Across Major SARS-CoV-2 Repositories. This table delineates the network attributes of various databases across distinct classifications: Authors, Country, Funder, and Institution. Metrics included are node count, edge count, clustering coefficients, and density scores.

|  |  | Number of Authors |  |  |  |  |  |  |  |  |  |
| --- | --- | --- | --- | --- | --- | --- | --- | --- | --- | --- | --- |
|  |  | 1 | 2-5 | 5-10 | 10-20 | 20-50 | 50-100 | 100-250 | 250-500 | 500-1000 | 1000 - 5000 |
| Repo | Year | Number of Papers |  |  |  |  |  |  |  |  |  |
| GISAI D | 2020 | 64 | 609 | 548 | 511 | 275 | 25 | 4 | 0 | 3 | 1 |
| GISAI D | 2021 | 93 | 911 | 1062 | 1207 | 634 | 60 | 7 | 0 | 5 | 1 |
| GISAI D | 2022 | 95 | 916 | 1135 | 1254 | 672 | 45 | 12 | 0 | 2 | 1 |
| ENA | 2020 | 72 | 420 | 304 | 178 | 104 | 18 | 9 | 1 | 1 | 0 |
| ENA | 2021 | 275 | 1219 | 608 | 410 | 198 | 28 | 14 | 0 | 1 | 1 |
| ENA | 2022 | 407 | 1347 | 875 | 525 | 180 | 30 | 12 | 4 | 0 | 1 |
| NCBI | 2020 | 80 | 613 | 546 | 288 | 114 | 14 | 3 | 0 | 0 | 0 |
| NCBI | 2021 | 126 | 1032 | 965 | 542 | 209 | 27 | 5 | 0 | 3 | 0 |
| NCBI | 2022 | 113 | 1189 | 1258 | 718 | 210 | 21 | 4 | 0 | 0 | 0 |
| DDBJ | 2020 | 15 | 49 | 36 | 16 | 3 | 0 | 0 | 0 | 0 | 0 |
| DDBJ | 2021 | 10 | 22 | 19 | 17 | 1 | 0 | 0 | 0 | 0 | 0 |
| DDBJ | 2022 | 6 | 32 | 23 | 23 | 7 | 0 | 1 | 0 | 0 | 0 |

**Table 3:** Number of Papers Per Number of Authors Across Major SARS-CoV-2 Repositories. This table displays the cumulative number of papers per number of author for each repository between 2020 and 2022.

| Database | Type | Clustering coefficient | Density | Mean degree | Mean weighted degree | Max community size | Mean community size | Median community size | Min community size | Total communities | Number of edges | Num of nodes |
| --- | --- | --- | --- | --- | --- | --- | --- | --- | --- | --- | --- | --- |
| DDBJ | Authors | 0.96 | 0.01 | 16.13 | 17.54 | 101.00 | 8.75 | 7.00 | 2.00 | 246.00 | 17359.00 | 2152 |
|  | Country | 0.42 | 0.08 | 4.56 | 16.70 | 43.00 | 19.00 | 12.00 | 2.00 | 3.00 | 130.00 | 57.00 |
|  | Funder | 0.77 | 0.68 | 8.15 | 44.15 | 13.00 | 13.00 | 13.00 | 13.00 | 1.00 | 53.00 | 13.00 |
|  | Institution | 0.60 | 0.01 | 5.24 | 5.60 | 71.00 | 4.80 | 3.00 | 2.00 | 87.00 | 1095.00 | 418.00 |
| ENA | Authors | 0.99 | 0.00 | 246.25 | 252.47 | 2987.00 | 11.28 | 6.00 | 2.00 | 6179.00 | 8583776.00 | 69,710 |
|  | Country | 0.71 | 0.29 | 43.54 | 9220.42 | 153.00 | 153.00 | 153.00 | 153.00 | 1.00 | 3331.00 | 153.00 |
|  | Funder | 0.93 | 0.92 | 14.71 | 1752.82 | 17.00 | 17.00 | 17.00 | 17.00 | 1.00 | 125.00 | 17.00 |
|  | Institution | 0.90 | 0.03 | 221.45 | 242.59 | 6395.00 | 68.30 | 2.00 | 2.00 | 101.00 | 763790.00 | 6898 |
| GISAID | Authors | 0.91 | 0.00 | 158.61 | 207.15 | 3310.00 | 18.25 | 8.00 | 2.00 | 4679.00 | 6771757.00 | 85,380 |
|  | Country | 0.65 | 0.26 | 40.50 | 823.06 | 159.00 | 159.00 | 159.00 | 159.00 | 1.00 | 3220.00 | 159.00 |
|  | Funder | 0.93 | 0.91 | 13.63 | 2289.00 | 16.00 | 16.00 | 16.00 | 16.00 | 1.00 | 109.00 | 16.00 |
|  | Institution | 0.31 | 0.00 | 27.35 | 48.77 | 4447.00 | 48.24 | 3.00 | 2.00 | 125.00 | 82,446.00 | 6030 |
| NCBI | Authors | 0.97 | 0.00 | 57.73 | 61.32 | 1088.00 | 10.59 | 7.00 | 2.00 | 7002.00 | 2140401.00 | 74,150 |
|  | Country | 0.56 | 0.19 | 27.31 | 741.80 | 148.00 | 148.00 | 148.00 | 148.00 | 1.00 | 2021.00 | 148.00 |
|  | Funder | 0.89 | 0.84 | 13.41 | 1448.12 | 17.00 | 17.00 | 17.00 | 17.00 | 1.00 | 114.00 | 17.00 |
|  | Institution | 0.30 | 0.00 | 24.98 | 32.79 | 5956.00 | 49.50 | 2.00 | 2.00 | 131.00 | 81002.00 | 6485 |

**Table 4.**
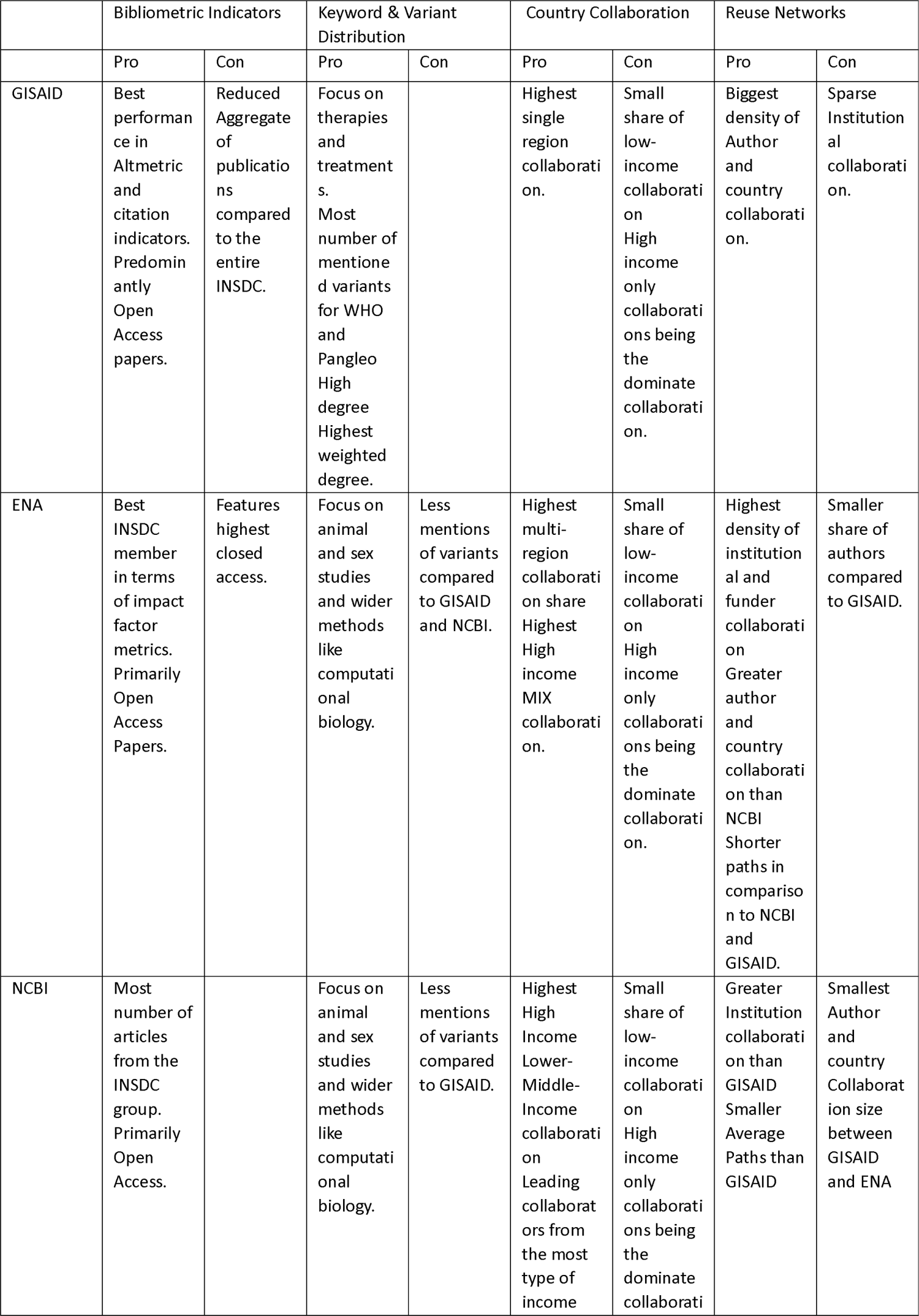
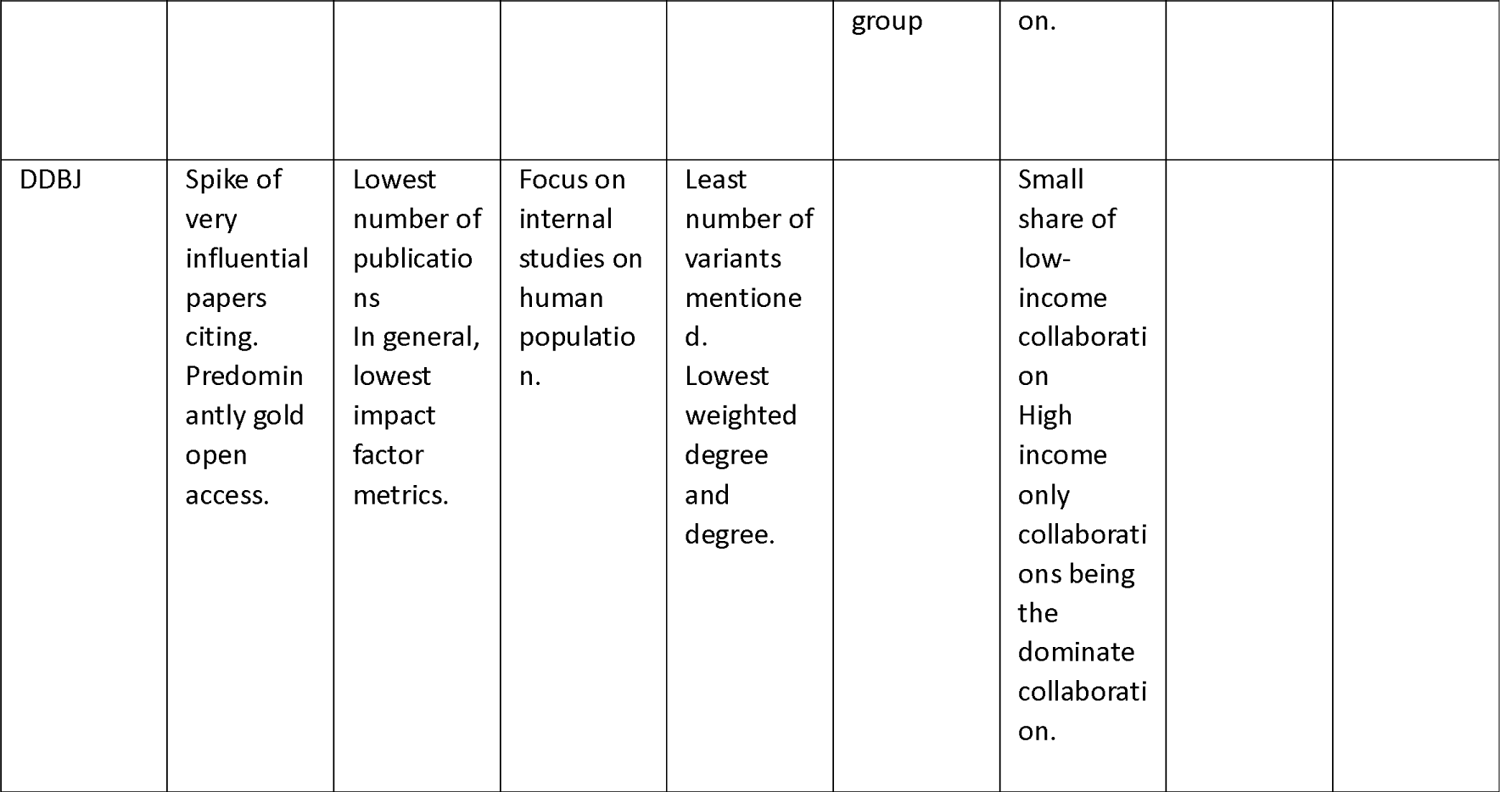
List of Pros and Cons for the strategies adopted by each repository based on results.

The country network also exhibited considerable homogeneity, albeit with more distinctive characteristics relative to the funding network. GISAID’s network emerged as the most expansive, boasting 159 nodes and 3220 edges. Most databases trended towards a singular community size, with DDBJ being an outlier with three communities. The apex clustering coefficients and density metrics were identified in the ENA network, registering at 0.71 and 0.29, respectively. The mean degree was highest for ENA at 43.54, while GISAID closely followed with 40.50, and NCBI and DDBJ were relatively lower at 27.31 and 4.56, respectively.

The institutional network landscape revealed intriguing variances. Notably, the ENA dominated in terms of nodes (6898) and edges (763790). The subsequent rankings were NCBI and GISAID with 6485 nodes (81002 edges) and 6030 nodes (82466 edges) respectively, while DDBJ lagged considerably at 418 nodes and 1095 edges. All databases manifested exceedingly low-density metrics, with ENA leading at 0.3. The clustering coefficients mirrored this trend, with ENA outpacing GISAID and NCBI by a factor of three and DDBJ by two. An interesting revelation was NCBI’s network comprising the largest number of communities at 131, superseding GISAID, ENA, and DDBJ, which had 125, 101, and 87 communities, respectively.

The “Authors” category emerged as the predominant network across databases. GISAID’s network was the most extensive with 85,387 nodes and 6,771,757 edges. NCBI and ENA followed with 74,155 nodes (2,140,401 edges) and 69,717 nodes (8,583,776 edges) respectively, while DDBJ was significantly smaller with 21,152 nodes and 17,359 edges. Owing to their extremely large size, density metrics were exceedingly low across all databases, with DDBJ being a notable exception. The clustering coefficient was predominantly high, with ENA leading at 0.99, trailed by NCBI and DDBJ at 0.97 and 0.96, respectively. In terms of community counts, NCBI was predominant with 70,002, followed by ENA, GISAID, and DDBJ with 6,179, 4,679, and 246 respectively. The community sizes (mean, median, and minimum) across databases exhibited remarkable consistency.

For the ENA database, in terms of connectivity degree of the network, the leading countries were as follows: the United States (126), the United Kingdom (124), Germany (112), Canada (105), South Africa (103), France (102), India (102), Australia (100), Brazil (97), the Netherlands (96), and China (95). In relation to betweenness centrality, the United States stood out with a score of 758.18. This was followed by the United Kingdom (652.73), France (510.59), Canada (455.33), and South Africa (322.51).

In the GISAID database, the countries with the highest degrees of connection were the United States (125), the United Kingdom (126), France (109), Germany (108), Brazil (104), Switzerland (103), South Africa (101), Canada (96), Saudi Arabia (95), Japan (94), and Egypt (93). When evaluating betweenness centrality, the leading countries were Senegal (653.05), Sweden (601.17), Italy (561.47), Australia (541.90), Brazil (474.20), Canada (411.64) and Nigeria (377.70).

For the NCBI database, the foremost countries in terms of degree were the United States (118), the United Kingdom (108), Germany (97), France (86), Italy (83), India (82), Australia (81), China (79), Switzerland (77), Canada (73), and Spain (72). Regarding betweenness centrality, France held the highest score with 708.41, followed by Italy (711.71), South Africa (563.36), the United Kingdom (495.84), and Portugal (469.37). For the DDBJ the leading countries in terms of degree were Japan (20), United Kingdom (19), United States (19), Italy (11), Sweden (10), Germany (10), Switzerland (10), Israel (9), Australia (9) and Croatia (8). Japan had the leading betweenness centrality score followed by United Kingdom, United States, China, Israel, and Australia.

In the DDBJ network, the predominant collaboration was between Japan and the United States, registering 94 instances, representing a significant majority. Subsequent collaborations included the United Kingdom and the United States (34), Australia and the United States (30), Japan and Norway (14), and Lebanon and the United Kingdom (14). Within the ENA database the largest global reach of country collaboration was recorded, with collaborations involving the United States dominated the top seven positions with China (33,298), Germany (22,125), the United Kingdom (20,314), Spain (16,911), Italy (16,750), France (14,899), and Japan (14,371). In the GISAID database, the United States also led country collaborations. But the top collaborative instances were far less with the United Kingdom (3,393), Germany (2,745), China (2,021), Canada (1,680), Spain (1,599), Australia (1,373), Japan (862), and India (835). A similar trend was identified in the NCBI database, where the United States featured prominently in the top eight collaborative positions. Collaborative instances included the United Kingdom (3,393), Germany (2,745), China (2,021), Canada (1,680), Spain (1,599), Australia (1,373), Japan (862), and India (835).

### Countries are color-coded based on their total number of publications

In the analysis of funding networks in figure 10, the principal funders for GISAID included ICRP, NIH, cOAlitionS, NSF, UKRI, DoD, and US Federal funders, exhibiting fifteen degrees of connection. Of these, UKRI demonstrated the most significant betweenness centrality. However, when considering the entire network, USDA held the highest betweenness centrality, valued at 57.5, with 12 degrees. The most substantial collaborations were observed between ICRP and US Federal Funders (2166 instances), followed by ICRP-NIH (2094) and NIH-US Federal Funders (2094). Subsequent significant collaborations were identified between cOAlitionS and ICRP (1433), EC & ERC (940), and UKRI (695). Within the NCBI network, the top funders in terms of connection degrees, totalling sixteen, were IRCP, NIH, cOAlitionS, UKRI, and US Federal Funders. In this group, UKRI retained the highest betweenness centrality, registering 32.5. Overall, NOAA exhibited the leading betweenness centrality at 68.67 with ten degrees. The primary collaborations were between ICRP and US Federal Funders (1755), NIH and US Federal Funders (1723), and ICRP and NIH (1720). Further collaborations included cOAlitionS with ICRP (711), EC & ERC (535), and US Federal Funders (390). For the ENA funding network, the dominant funders in terms of connection degrees, all at sixteen, were ICRP, NIG, cOAlitionS, AMRC, EC & ERC, NSF, UKRI, HRA, CDC, and US Federal Funders. Among these, CDC displayed the highest betweenness centrality at 48.67, but in a broader context, NOAA with twelve degrees possessed a greater betweenness centrality of 67. The chief collaborations were with ICRP and US Federal Funders (1374), NIH and US Federal Funders (1317), and ICRP and NIH (1314). This was followed by collaborations involving cOAlitionS and ICRP (1113), EC & ERC (940), and UKRI (740). In the DDBJ network, the US Federal Funders emerged as the primary funding group, reflecting fourteen degrees of connection. The maximum betweenness centrality was attributed to NSF, although it only had ten degrees. Predominant collaborations were discerned between ICRP and US Federal Funders (44), ICRP and NIH (43), and NIH and US Federal Funders (43), followed by CDC and US Federal Funders (24). Additional collaborations involved cOAlitionS and ICRP (23), EC & ERC (19), US Federal Funders (14), and UKRI (12).

In the institutional network presented in figure 11 the University of Oxford (826), followed by Imperial College London (668), University of Edinburgh (651), Harvard University (638), the Ministry of Health (596), University of Washington (570), University of Cambridge (537), University College London (504), London School of Hygiene and Tropical Medicine (501) and the Institut Pasteur (462) were the leading nodes for GISAID. In the same network, the University of Oxford again leads in the largest betweenness centrality with 604373.83, followed by Harvard University, University of Hong Kong, University of Edinburgh and the Ministry of Health with 564449.4, 478585.8, 435489.2 and 405605.17 respectively. The top five leading collaborations were between the Fred Hutchinson Cancer Centre and the University of Washington (141), KU Leuven and Rega Institute for Medical Research (116), Imperial College London and the University of Oxford (113), Howard Hughes Medical Institute and University of Washington (73) and the Institute of Microbiology and the University of Chinese Academy Science (73).

In contrast, the ENA platform showcases a broader scope of institutional collaborations. The Sun-Yat Institute (also referred to as Zhongshan University) emerges as the most connected with 2593 degree, followed by the University of Paris (2361), University of Oxford (2031), University College London (1987), Imperial College London (1894), Harvard University (1876), University of Cambridge (1811), University of Edinburgh (1738), Kings College London (1732) and Stanford University (1695). Institutions pivotal in the genesis of ENA, EMBL-EBI, namely the European Bioinformatics Institute and the European Molecular Biology Laboratory, also feature prominently, with the European Bioinformatics Institute receiving 1637 degree and the European Molecular Biology Laboratory with 1542. The leading institutes in terms of betweenness centrality are University College London, University of Oxford, University of Cambridge, Harvard University, and Sun Yat with 366428.6, 363747.68, 258835.77, 244949.02, 242804.76 respectively. The leading collaboration between institutes were the British Medical Association and the Canadian Medical Association (80), followed by the University of Cambridge and the Wellcome Sanger Institute (75), Broad Institute and Harvard University (69), Imperial College London and University of Oxford (61) and the European Molecular Biology Laboratory and Heidelberg University (59). The European Molecular Biology Laboratory also had high collaborations with the German Cancer Research Center (51), while the European Bioinformatic Institute had substantial collaborations with the University of Cambridge (45), the Wellcome Sanger Institute (44) and University College London (37).

For the NCBI, the leading institutions in terms of degree were Harvard University (820), University of Cambridge (693), Wellcome Sanger Institute (498), University of Washington (496), Cornell University (487), Stanford University (477), University College London (476), National Center for Biotechnology Information (475) and the University of Melbourne (453). In terms of betweenness centrality, Harvard University overshadowed the rest with a score of 724458.49, followed by University of Oxford, University of Melbourne, Imperial College London and the University of California with 519537.68, 346122.59, 316106.06, 213986.81 respectively. The leading institutions in terms of collaboration were the BGI Group (China) and the University of Chinese Academy of Science (80), Broad Institute and Harvard University (58), Harvard University and Massachusetts General Hospital (39), Brigham and Women’s Hospital and Harvard University (37) and Broad Institute and Massachusetts Institute of Technology. In the top twenty of collaborations, the NCBI themselves did not appear.

For the DDBJ, the leading institutes in terms of degree were Harvard University (49), National Institute of Infectious Diseases (48), University of Tokyo (32), Kyoto University (30), Broad Institute (26), Waseda University (24), Hokkaido University (23), Keio University (22), European Bioinformatics Institute (22) and Cornell University (22). In terms of betweenness centrality, Harvard, National Institute of Infectious Diseases, European Bioinformatics Institute, Kyoto University and Research Organization of Information and Systems with 6304.96, 4225.42, 3541.08, 2943.33 and 2851.85 respectively.

## 5 Discussion

A significant distinction between the two data governance models is observed in the variance of MeSH (Medical Subject Headings) keywords and the distribution of variants. GISAID possesses the most significant proportion of variant names within its corpus, lending credence to the notion that its primary objective is to monitor novel variants, assess genetic mutations, and evaluate vaccine immunity in relation to these variations. This is corroborated by the relative distribution of keywords in which GISAID shows a marked inclination towards public health applications and therapeutic-centric terminologies, while INSDC captures a wider spectrum of biological and epidemiological research domains, including substantive efforts in upstream research. Such distinctions manifest in the kind of data each model disseminates: GISAID predominantly shares RNA or protein data, which it provides in a ready-to-use format for immediate employment by public health officials, whereas the INSDC provides a broader spectrum of epidemiological data, encompassing a considerable quantity of raw reads and annotation data and providing more opportunities for linking the data to other existing resources.

This difference in audiences and usages goes some way towards account for the difference in emphasis in the governance models preferred by the two infrastructures. While GISAID depends on the comprehensiveness of its respondents and participants to be able to provide as wide-ranging a picture of global mutation patters as possible, INSDC is more focused on facilitating linkage between datasets, thereby supporting discovery on a variety of novel aspects of SARS-CoV-2 behaviour and interactions with host organisms and environments. Openness in the sense of immediate, wide-ranging usability of data is therefore arguably more crucial to INDSC, while attention to which sources are captured and the extent to which they can represent the world-wide situation is of primary concern for GISAID.

The disproportionate size of the DDBJ corpus compared to NCBI or ENA can be understood in the context of a government mandate by the Ministry of Health, Labour and Welfare in Japan which required all SARS-CoV-2 genomes to be registered with GISAID, not elsewhere (MHLW, 2021). In turn, this meant sequences submitted to GISAID from Japan were not allowed to go into an INSDC repository due to licencing terms.

Even when taking this difference in emphasis into account, the results from the analysis above presents a mixture of benefits and drawbacks to GISAID and the members of the INSDC in their capacity to support diverse research collaboration for the study of SARS-CoV-2, and thus deserves some critical reflection and contextualization.

For a start, during the 2020-2022 pandemic years, the GISAID corpus has witnessed an increased frequency of publication, citation, and discussion compared to other INSDC members. This underscores GISAID’s instrumental role in forging early collaborative efforts between global researchers and institutions (Khan et al., 2021). However, when combining the output of all INSDC members, the cumulative publications significantly eclipse that of GISAID and demonstrate comparable citations and Altmetric scores. While there are indications of GISAID’s publication volume diminishing in 2023, both the ENA and NCBI are observing an upward trajectory in their publication count. The slow start exhibited by INSDC members may be attributable, in part, to the delayed release of COVID-19-specific data portals and services, like CV19DP and the NCBI SARS-CoV-2 Data Dashboard, until 2021.

Since their inception, these platforms have undergone iterative enhancements, encompassing new functionalities for data submission, retrieval, and linkage (Rahman et al., 2023). Nevertheless, this growth should be approached circumspectly. The rising prevalence of both gold and green access classifications, coupled with the emergence of hyper authorship papers, comprising 50+ authors within both GISAID and INSDC, illuminates a trajectory towards augmented collaboration and transparency in virological inquiries. At the same time, this highlights the conundrum where fully open repositories like ENA may simultaneously amplify the frequency of restricted access and solo authorship articles at the same time. This raises the question of whether the unrestricted data sharing paradigm, endorsed by INSDC members, inadvertently leans towards an object-oriented perspective on Open Science, where the sheer volume of outputs being shared may overshadow diversity and inclusivity (Leonelli 2023).

Beyond these emerging concerns around the meaning and implications of openness for research, our findings also point to the potential dangers of implementing openness in ways that inadvertently constructs new obstacles in the way of scientific collaboration. Both GISAID and INSDC exhibit high representation of data from high-income countries. Adding to this, the country collaboration networks show there is a distinct prominence of certain nations in the global bioinformatics landscape. The United States and the United Kingdom recurrently emerge as frontrunners in terms of connectivity degree across these platforms situating them as key actors in the genomic surveillance landscape. Recurrent actors weren’t only limited to countries. The heterogeneity observed between the funding network for GISAID and INSDC, for instance, raises important questions about the extent to which funding sources influence the distribution and accessibility of research resources. The results similarly show institutional hierarchies and clustered collaborations across GISAID, ENA, NCBI, and DDBJ platforms. Unsurprisingly, esteemed institutions like the University of Oxford, Harvard University, and University College London consistently appear pivotal, either by connection degrees or betweenness centrality. Moreso, the general dominance of inter-regional partnerships- or in the case of ENA intra-regional connections mainly with high income countries – in research collaborations raises questions about the extent to which research data accumulated in centralised data infrastructures can truly be considered a global endeavour, or whether it is still largely driven by a select few countries, institutions, income groups and funders.

These results raise concerns about the representation of data from low-income countries within both data governance models, which raises important questions about access to and sharing of scientific resources, as well as the potential for biases in data sampling based on incomplete data – which align with emerging scholarship questioning the accessibility and equitable distribution of supposedly open scientific resources (UNESCO Recommendations 2021, Ross-Hellauer et al 2022, Leonelli 2023). Efforts to improve data sharing and promote equity in scientific research are critical for ensuring that all populations, regardless of geography or economic status, have access to the best available information and resources for preventing and treating disease (Prat and Bull 2021; Cousins et al. 2021; Staunton et al. 2021).

Characteristics of the lesser prominent collaborations between each repository offer revealing insights. For instance, in the GISAID country network Kenya, Nigeria, Senegal and South Africa make up a decent share of degree and betweenness centrality. This united front may in part be due to the federated effort by African nations to share data between themselves using GISAID to keep their data secure (Tegally et al., 2022). This might suggest the GISAID model has been more effective in building trustworthiness among users from different resource environments to collaborate and engage in data generation (ODI, 2019; RDA, 2020). It also seems to reinforce claims by Bernasconi (2021) that a partially closed access model is preferred at a global scale for viral data sharing. In contrast to Bernasconi, however, our results also suggest that GISAID’s network topology appears less densely interconnected than that of the ENA or NCBI. ENA’s pronounced interconnectivity might suggest either the emergence of specialized research clusters or a propensity towards insular data exploration as evidenced by their large growth in single author papers. Interestingly, data trends support the latter option, with ENA fostering broader multi-regional collaborations, potentially benefiting from its commitment to data interoperability. Placing emphasis on interoperability and usability by members of the INSDC may permits users of the infrastructure to explore their own research questions and methods more easily by linking together the wider variety of research data types. However, GISAID’s low density and high average path may positively reflect the controlled access to the repositories data and the repositories tendency to have much larger authorship per paper. These results warrant further exploration into the communities and segregated communities’ part of each repository.

## 6 Conclusion

Our bibliometric analysis highlights the strengths and limitations of the regulated and unrestricted approaches to open data governance embodied by GISAID and INSDC, respectively. These databases furnish valuable resources for scientific research, yet they diverge in their bibliometric indicators, country and income collaboration and corpus networks. While data sharing initiatives advocating for complete openness like members of the INSDC highlight the advantages of immediate data access, they tend to overlook the sociocultural, institutional, and infrastructural factors that affect data reuse. These factors include disparities in geo-political locations, power dynamics among research sites, expectations regarding intellectual property, funding availability, and digital connectivity resources. GISAID’s partially open model offers an alternative approach that has drawn more diverse geo-political locations into its fold. However, despite this increased geographical representation, it does not necessarily translate into greater epistemic diversity within research topics. This can limit the breadth of scientific perspectives and inadvertently funnel research into narrow or pre-established trajectories. While the different user base of GISAID and INSDC can explain some of the discrepancies between these two systems, their respective approaches to openness arguably account for at least some of the attitudes and preferences of their users, including the greater emphasis on wide-ranging and exploratory biological research facilitated by INSDC.

Looking ahead, while our current analysis has placed significant emphasis on betweenness centrality measurements, future research could explore community segregation between institutions, thereby uncovering deeper insights into the inherent collaborative or insular behaviours of academic institutions in the context of data sharing. The landscape of data governance is rife with contention, especially concerning what constitutes as responsible and ethical practices. Yet, both GISAID and INDSC, in INSDC respective ways, demonstrate and support effective modes of collaboration. Through critical reviews of such data repositories, we can ascertain the nuances and implications of their governing structures. Through empirical exploration and better understanding of these intricacies, the academic community stands a better chance to design robust and inclusive systems for data governance that truly foster global scientific collaboration.

## Acknowledgements

We thank colleagues at the European Bioinformatics Institute, the “Philosophy of Open Science for Diverse Research Environments” (PHIL_OS) project and the Exeter Centre for the Study of the Life Sciences (Egenis) for their feedback and support in this research, as well as Carole Goble for illuminating discussions and the attentive audience of the PhilInBioMed conference in Pittsburgh (November 2022) where this work was first presented.

## Footnotes

1 Note that our discussion is not meant to be comprehensive of all possible forms of data governance in this domain. Rather, we take these two cases as exemplifying significant and widespread models of data sharing, which are worth comparing to further enhance existing understandings of best data management practice.

2 Given these controversies, this paper does not aim to take a strong position on whether GISAID in fact complies with the FAIR principles, which would require a different kind of analysis and empirical evidence (including checks on the extent to which GISAID data have been accessible in practice). Rather, we focus on the GISAID governance mode as articulated by the infrastructure itself, according to which data are Accessible upon request and compliance with the GISAID license agreement; Findable on the GISAID database; Interoperable as long as users declare prospective purposes and delimit the degree to which GISAID data are integrated with other data sources; and Reusable as long as the provenance of data is clearly acknowledged and the prospective use serves public health goals.

